# Systems biology framework for rational design of operational conditions for *in vitro / in vivo* translation of microphysiological systems

**DOI:** 10.1101/2025.01.17.633624

**Authors:** Jose L. Cadavid, Nikolaos Meimetis, Tyler Matsuzaki, Erin N. Tevonian, Linda G. Griffith, Douglas A. Lauffenburger

**Author notes:** These authors contributed equally (Jose L. Cadavid, Nikolaos Meimetis).

## Abstract

Preclinical models are used extensively to study diseases and potential therapeutic treatments. Complex in vitro platforms incorporating human cellular components, known as microphysiological systems (MPS), can model cellular and microenvironmental features of diseased tissues. However, determining experimental conditions -- particularly biomolecular cues such as growth factors, cytokines, and matrix proteins -- providing the most effective translatability of MPS-generated information to in vivo human subject contexts is a major challenge. Here, using metabolic dysfunction-associated fatty liver disease (MAFLD) studied using the CNBio PhysioMimix as a case study, we developed a machine learning framework called Latent In Vitro to In Vivo Translation (LIV2TRANS) to ascertain how MPS data map to in vivo data, first sharpening translation insights and consequently elucidating experimental conditions that can further enhance translation capability. Our findings in this case study highlight TGFβ as a crucial cue for MPS translatability and indicate that adding JAK-STAT pathway perturbations via interferon stimuli could increase the predictive performance of this MPS in MAFLD studies. Finally, we developed an optimization approach that identified androgen and EGFR signaling as key for maximizing the capacity of this MPS to capture in vivo human biological information germane to MAFLD. More broadly, this work establishes a mathematically principled approach for identifying experimental conditions most beneficially capturing in vivo human-relevant molecular pathways and processes, generalizable to preclinical studies for a wide range of diseases and potential treatments.

## 1. Introduction

Ethical, practical, and economic constraints on human subject research for the study of mechanisms of disease and their treatment necessitate the use of preclinical animal models of disease. While animal models are a gold standard in preclinical research, they have only an approximately 10% rate, broadly, of successful translation to the clinic^1^ and often fail to predict drug toxicities due to inherent inter-species differences^2^. An emerging alternative approach is that of *in vitro* complex cell culture models engineered with human cells, also known as microphysiological systems (MPS), as they can avoid inter-species differences. MPS platforms can be designed to comprise diverse cell types, extracellular matrix components, and diffusible molecular stimuli relevant to a particular disease. Not surprisingly, this approach is being increasingly adopted by industry^3,4^ and regulatory agencies^5^. However, MPS exist along a wide spectrum of biological and operational complexity associated with a range of experimental throughput and cost. Therefore, defining biological parameters (cell types, extracellular matrix components, molecular stimuli), or ‘cues’, for the operation of a given MPS to enhance its translational potential is crucial for their wider adoption as useful preclinical models.

Among disease areas for which MPS platforms are being used extensively is metabolic dysfunction–associated fatty liver disease (MAFLD)^6–8^ and hepatotoxicity^9,10^. Non-alcoholic fatty liver disease (NAFLD), a form of MAFLD, is one of the most common liver diseases in the world, occurring in almost 25% of the population worldwide^11^. MAFLD’s clinical outcome ranges from simple steatosis to metabolic dysfunction-associated steatohepatitis (MASH; previously called NASH) and can lead to liver cirrhosis^12^. Fat-tissue-derived free fatty acids (FFAs), among other products, can lead to steatosis and lobular inflammation. Accumulation of inflammatory-associated cells, cytokines, and other signals can lead to chronic necroinflammation, marking an MAFL-to-MASH transition. Chronic hepatocyte cell death and mild to advanced fibrosis, together with increased levels of TNFs, TGFβ, and IL-18, result in increased hepatocellular carcinoma (HCC) risk^12^. The complexity of the disease and the plethora of molecular signals in the tissue microenvironment have led to the development of a wide array of *in vitro* models of disease that mimic specific phenotypes of interest, such as fibrosis and steatosis, each providing different trade-offs between throughput, complexity, and translatability^13^.

Absent a complete understanding of the molecular and cellular processes driving disease pathophysiology, the definition of ‘cues’ required *in vitro* to achieve good MPS translation is often based on putative factors based on prior knowledge of disease biology^4^. Direct functional and molecular (e.g. transcriptomic) comparison between *in vitro* and *in vivo* systems^14,15^ can be used to rank MPS models for specific contexts of use^16^, and computational comparison at the pathway level can be used to assess whether a perturbation-induced response of an *in vitro* model is similar to the response observed *in vivo*^17^. However, these approaches assume the need for a direct one-to-one mapping between MPS and humans (Figure 1a) and do not inherently approximate an alternative function that can link the *in vitro* and *in vivo* biological space. Moreover, even simple molecular cues provided to monocultures in a 2D MPS can result in clinically relevant responses to perturbations and treatments^18,19^ without the need to increase model complexity. Hence, a robust framework is necessary for comparing *in vitro* and *in vivo* data to determine a set of informative cues that can increase the predictive power of MPS models without necessarily increasing MPS complexity.

**Figure 1:**
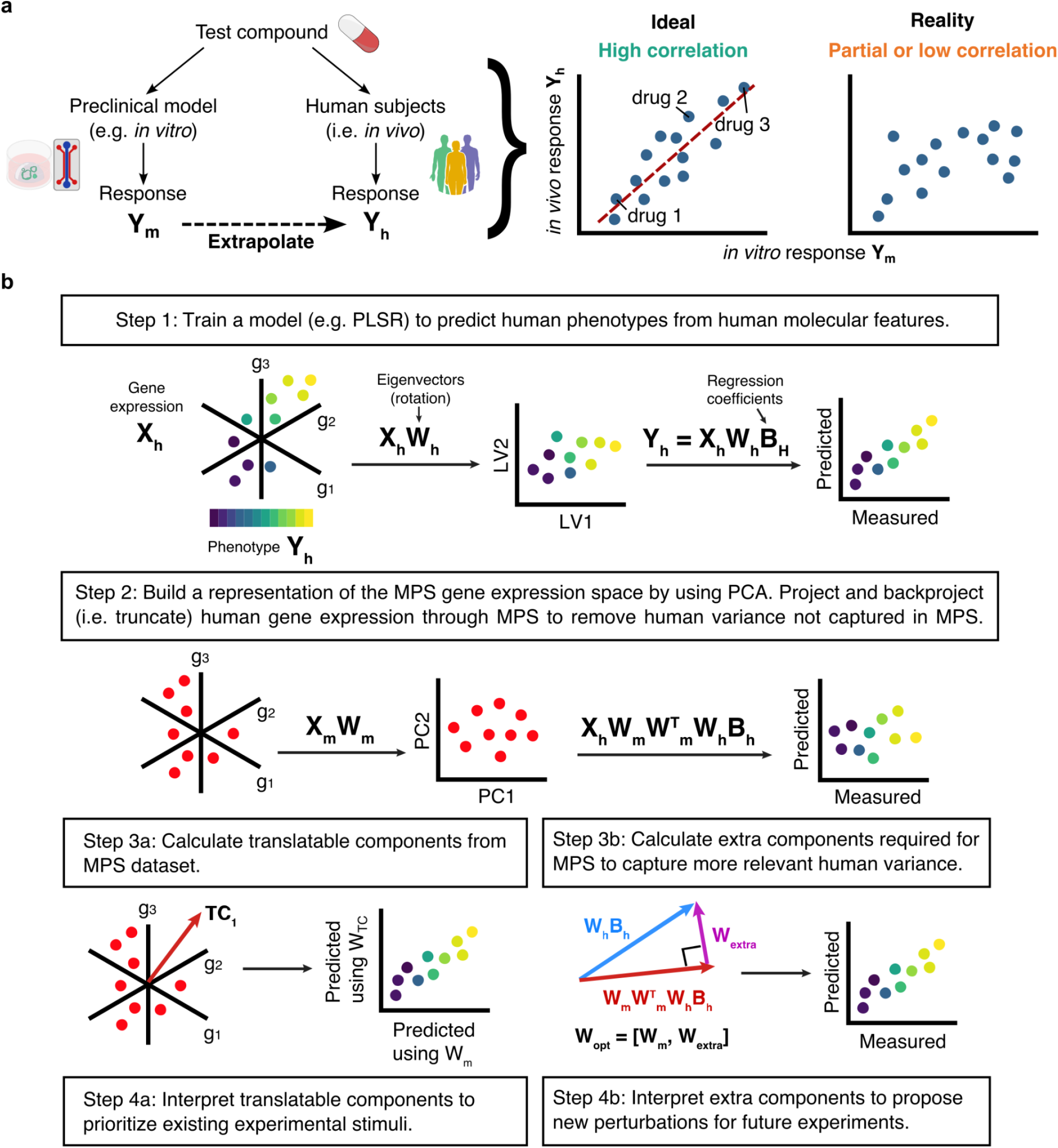
Data-driven modeling pipeline to identify translatable and missing information from microphysiological systems and human data. **(a)** Microphysiological systems (MPS) and other preclinical models are designed to predict the response of human subjects to a particular perturbation, such as a drug or novel compound. In the ideal case, the response to perturbations of the MPS would correlate well with human subjects. However, in practice, such correlations might be low or only partial when assuming one-to-one mapping between systems. **(b)** Our approach LIV2TRANS (Latent *In Vitro* to *In Vivo* Translation) finds translatable information between *in vitro* and *in vivo* systems by operating in a latent space built from molecular features measured in both systems.

A data-driven systems biology framework, in combination with Machine Learning (ML) computational modeling, can provide the tools to enable the systematic understanding and design of *in vitro* models^20^. Systems biology as a field has enabled studying complex biological systems by observing their response under multiple conditions^21,22^ and developing tools that allow comparing different biological systems (e.g. *in vitro* and *in vivo* models). Additionally, ML can fill this gap by acting as a global function estimation framework that maps observations between *in vitro* and *in vivo* models. Existing work has focused on either translating the molecular profiles between different biological systems^23–25^ or making clinically relevant predictions about stimuli observed in experimental models, identifying in the process molecular features of the reference experimental system (typically animal models) that are most germane for understanding phenotypes in another system (typically human subjects)^26–29^. These approaches enable the translation of observations made in the experimental models back to the humans in the clinic, but they are not inherently designed to identify conditions to enhance an existing experimental system for better recapitulating the observed human phenotypic characteristics.

In our new study here, using MAFLD as a case study, we developed an ML approach for the rational design of cues to be added to an *in vitro* system. Specifically, our approach leverages molecular and phenotypic data from human subjects^30^ and an MPS model^8^ to summarize molecular signatures into latent variables and decompose disease phenotypes (e.g. liver fibrosis), enabling their prediction. The latent variables can inform the selection of cues along a newly identified molecular direction, optimizing the MPS model to capture more of the phenotypically relevant molecular variability displayed in humans. Importantly, rather than aiming to replicate human MAFLD/MASH precisely, our approach leverages the controllability of MPS models to propose cues that can facilitate better *in vitro*/*in vivo* translation. Interrogation of the molecular contributions along this patient-pathology axis, via pathway activity inference, reveals that perturbation of the JAK-STAT signaling pathway via interferon stimulation can enhance the predictive power of MPS by generating disease-relevant biological variance *in vitro*. This increased predictive power using this augmented molecular space is generalizable to another two clinical human datasets^31,32^, which were used as external test sets for the validation of the approach. Moreover, the observed JAK-STAT signaling perturbation can be identified when repeating the analysis for other pairs of *in vitro*^6,7^ and clinical datasets^31,32^. Finally, recognizing that *in vitro* models should also capture broad human biological variance, we used an optimization algorithm to widen the total captured variance by the MPS and observed that, while not being the most dominant molecular direction proposed, perturbing JAK-STAT signaling can also increase the total captured human variance. We used publicly available transcriptomic datasets^33^ to identify stimuli that can perturb the MPS model along the identified molecular directions. Altogether, our approach provides a principled way for selecting cue conditions that align with relevant phenotypic directions in humans, thereby advancing our understanding of disease mechanisms and their therapeutic solutions.

## 2. Results

### 2.1. A model translation pipeline to identify translatable and missing information in microphysiological systems

To be informative, a preclinical model should capture the phenotypically relevant human variation of the clinical data. Ideally, an MPS would be designed in a way that allows one-to-one mapping between *in vitro* and *in vivo* conditions. However, such a requirement might be unnecessarily restrictive and would likely require the use of highly complex *in vitro* models. Moreover, there is no guarantee that a single *in vitro* model will correlate well with human data across all potential scenarios, and in reality, the mapping between *in vivo* and *in vitro* systems is only partial (Figure 1a). We accordingly employ a data-driven modeling framework that maps information from *in vitro* models to *in vivo* without explicitly requiring one-to-one mapping by exploiting the covariance among common predictive molecular features measured in both contexts, similarly to that recently demonstrated for improved translation of preclinical animal model data^34^.

On this front, we developed a Machine Learning (ML) approach named Latent *In Vitro* to *In Vivo* Translation (LIV2TRANS) (Figure 1b, Methods section) for the rational design of cues to identify translatable and missing information in *in vitro* systems, focusing on an MPS model of MAFLD/MASH as a case study. Our approach aims to link *in vitro* and *in vivo* systems through a set of molecular features (e.g. transcriptomics) that are predictive of human phenotypes and that are measured in both systems. Instead of assuming that particular molecular features should match between systems, or domains, we posit that a more important requirement for translation is that the MPS model should capture human-relevant biological variance to be informative. Practically, this means that if human gene expression is predictive of the human phenotype, then minimum predictive power should be lost when discarding biological variance not captured in the MPS system.

As a first step, we construct a model to predict human phenotypes from molecular data to determine its information content (Figure 1b-1). For our particular case study, we used partial least squares regression (PLSR) to build a model because it accommodates more than one predicted phenotype, and it calculates latent variables that contain phenotype-relevant variance in the data. We then build a representation of the gene expression space spanned by the MPS model using principal component analysis (PCA), and project human gene expression data onto this MPS-PC space (Figure 1b-2). This projection truncates human data to the gene expression space spanned by the MPS, and therefore eliminates biological variance not captured *in vitro*. Projected human data are then back-projected to human gene expression space and used to predict human phenotypes with the model trained in step 1. The loss in predictive power of the model when using human data that has been “passed” through the MPS-PC space is a quantitative metric of the extent to which the MPS system captures human-relevant molecular variance. In what follows, we refer to the projection/back-projection step as “truncating” the human data for simplicity.

Generally, the MPS-PC space might only contain a limited number of directions in gene expression space that are relevant for preserving human-related biological variance. Thus, to determine these translatable components (TC), we find a subset of linear combinations of PCs that can be used instead of the full set of PCs for truncating human data without reducing the predictive power of the MPS model (Figure 1b-3). We then identify a set of extra latent variables, defined as a set of directions in gene expression space orthogonal to the MPS-PC space that would enable the retrieval of human-relevant biological variance lost during truncation of the human dataset through the original MPS dataset (Figure 1b-3). Interrogating the TCs can enable the identification of cues already present in the MPS that are key for capturing the human phenotype, while the extra latent variables can inform the selection of new cues that generate additional biological variance in the MPS to potentially capture more of the human-relevant molecular variability in future MPS experiments. Summarizing these mathematical steps conceptually, instead of aiming to replicate human phenotypes, our approach leverages the controllability of MPS models to propose cues that can facilitate better *in vitro*/*in vivo* translation by pooling the variance, hence information, contained in a set of MPS models with diverse cues.

### 2.2. Human gene expression is predictive of MAFLD/MASH histological scores

The first step in our computational pipeline was to build a mathematical model that links the measured human molecular features to the relevant human phenotype. Using MAFLD/MASH as a case study, we re-analyzed a large clinical dataset obtained from human biopsies collected across different stages of the disease and characterized with RNA-sequencing^30^. Since both the MAFLD score and fibrosis stage are clinically relevant for the diagnosis and prognosis of MAFLD/MASH^35,36^, we sought to model both phenotypes simultaneously from gene expression data. While it is possible to build independent models for each phenotype^32^, in reality, both scores correlate as the disease progresses (Figure 2b), and modeling them independently ignores the biology common to both. Therefore, we trained a partial least squares regression model (PLSR) to predict both phenotypes jointly. PLSR reduces the dimensionality of the gene expression data and projects the human samples onto a set of latent variables required to predict a set of phenotypes, and therefore provides an estimate of the gene expression subspace relevant to model MAFLD/MASH. The projected human scores are then regressed against each phenotype of interest, linking gene expression to phenotype. After tuning the optimal number of latent variables (see Methods and Supplementary Fig. 1), we trained PLSR models using a 10-fold cross-test split with all the data. The resulting PLSR model remains predictive in 10-fold cross-validation and performs significantly better than models built with shuffled data (Figure 2c) and generalizes well to independent clinical datasets (Supplementary Fig. 2). Having verified the robustness of the PLSR model, we re-trained the model with the complete dataset and obtained predictions that correlate well with the annotated scores (Figure 2d) within training data (Spearman’s rank correlation 0.85 and 0.94, respectively). Interestingly, the total variance explained by the eight latent variables in the model corresponds to only 42% of the total gene expression variance (around 15% in the first two latent variables; Figure 2e), suggesting that a significant portion of the variability in human gene expression data is not informative of the MAFLD/MASH stage. Overall, these results confirm that gene expression contains information about disease progression and indicate that the PLSR model captures the gene expression variance relevant to characterize the progression of human MAFLD/MASH.

**Figure 2:**
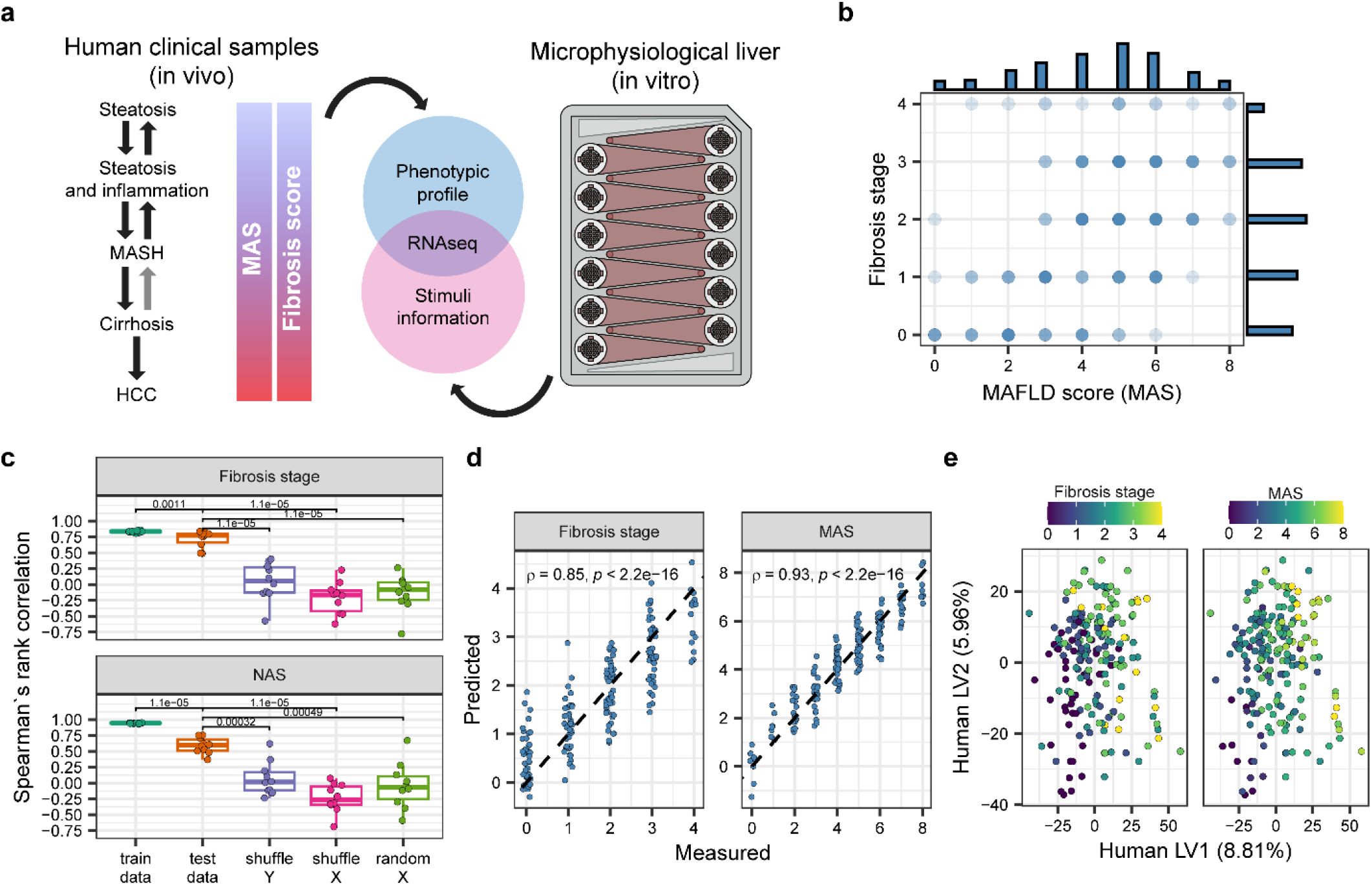
Modeling MAFLD/MASH phenotypic scores from bulk transcriptomics. **(a)** Bulk RNAseq data is used to translate between clinical samples (*in vivo*) with phenotypic annotations and a microphysiological model of disease (*in vitro*). **(b)** The MAFLD score (MAS) and Fibrosis stage are clinically relevant metrics of disease that correlate and need to be modeled jointly. **(c)** A partial least squares regression (PLSR) model predicts both phenotypes from bulk transcriptomics and performs significantly better in 10-fold cross-validation than random or shuffled models. For all comparisons, a two-sided unpaired Wilcoxon test was used. In all boxplots, the centerline denotes the median, the bounds of the box denote the 1st and 3rd quantiles, and the whiskers denote points not being further from the median than 1.5 × interquartile range (IQR). **(d)** The PLSR model with eight latent variables fits the full dataset well and achieves good correlations with the measured phenotypes. **(e)** Qualitative trends in the phenotypic scores are visible when projecting the human data onto the first two latent variables of the PLSR model.

### 2.3. Biological human variance predictive of MAFLD/MASH scores is not fully captured in a microphysiological model of disease

Having built a model to predict human MAFLD/MASH scores from gene expression, we next evaluated the extent to which a well-characterized liver MPS captures human-relevant biological variance (Figure 2a). We chose this particular MPS for our case study because it recapitulates key phenotypic features of MASH^8,37^. Further, the authors collected a large compendium of gene expression data obtained from multiple experimental conditions that included combinations of exogenous TGFβ, cholesterol, lipopolysaccharide (LPS), fructose, free fatty acids, and different ratios of hepatocytes and non-parenchymal cells (NPC)^8^. Using such MPS data was important since our approach relies on calculating common covariance structures between MPS and human subjects, which benefits from using large and comprehensive datasets.

As a first step, we sought to quantify the transcriptional similarity between the MPS samples and human samples. We scored the different MPS samples using a published gene expression signature that correlates well with MAFLD score and fibrosis stage^32^ (Supplementary Fig. 3a-3b). The particular combination of experimental cues added to the MPS had a significant impact on the calculated enrichment scores, with samples containing TGFβ displaying an overall higher enrichment score for both phenotypes. Unsurprisingly, this result is consistent with known disease biology since TGFβ is a strong fibrogenic factor known to be involved in the progression of MAFLD^38,39^ and other fibrotic diseases^40,41^, which is also why it was included in the original MPS study. However, further comparisons between each of the MPS and human samples revealed that most MPS samples correlated more strongly with human samples with intermediate fibrosis stages (1-3) and MAFLD scores (4-6) (Supplementary Fig. 3c-3d). This analysis highlighted that while the MPS model captures some disease-relevant biology, it does not map directly to the full range of human disease as assessed by direct gene expression similarity.

Hence, we used our data-driven approach (Figure 1) to map the MPS dataset to the human dataset in a more unbiased way that does not require one-to-one sample mapping. We performed principal component analysis (PCA) on the MPS dataset after averaging experimental replicates to preserve only gene expression variation due to changes in experimental treatments (Figure 3a). Since principal components (PCs) are the eigenvectors of the covariance matrix of the gene expression matrix, they span a gene expression subspace where the MPS has a non-zero variance. We projected the human gene expression onto the MPS PCA space and then back-projected the data to the original human space so that it could be projected onto the latent variables of the PLSR model (Figure 2e) to assess changes in the predictive performance of the model. The projection/back-projection step, or truncation, effectively removed human gene expression variance not captured in the MPS-PC space, as evidenced by a smaller spread of the human data when visualized in the latent variables of the PLSR model (Figure 3b). Moreover, truncation of the human data through the MPS gene expression space resulted in a significant drop in the predictive capability of the PLSR model, as shown by the lower correlation coefficients between the measured and predicted phenotypic scores (Figure 3c). Note that data truncation is mathematically equivalent to finding the best representation of human data as linear combinations of MPS samples (Supplementary Note). While retraining the regression coefficients in the PLSR model improved the apparent fit between measured and predicted scores, it did not increase the correlation coefficients to the values obtained with the non-truncated human dataset when assessed in individual data partitions (Supplementary Fig. 4). This indicates that the truncation step removed important biological information necessary for resolving differences between human phenotypes. Finally, we validated that the drop in predictive performance due to the truncation is not an artifact of the PLSR modeling by comparing the performance before and after truncation with other machine learning and deep learning algorithms (Supplementary Fig. 5). All together, these results suggest that the MPS dataset used in this case study does not capture all of the important disease-relevant human variance, even if it recapitulates certain known features of MASH^8^.

**Figure 3:**
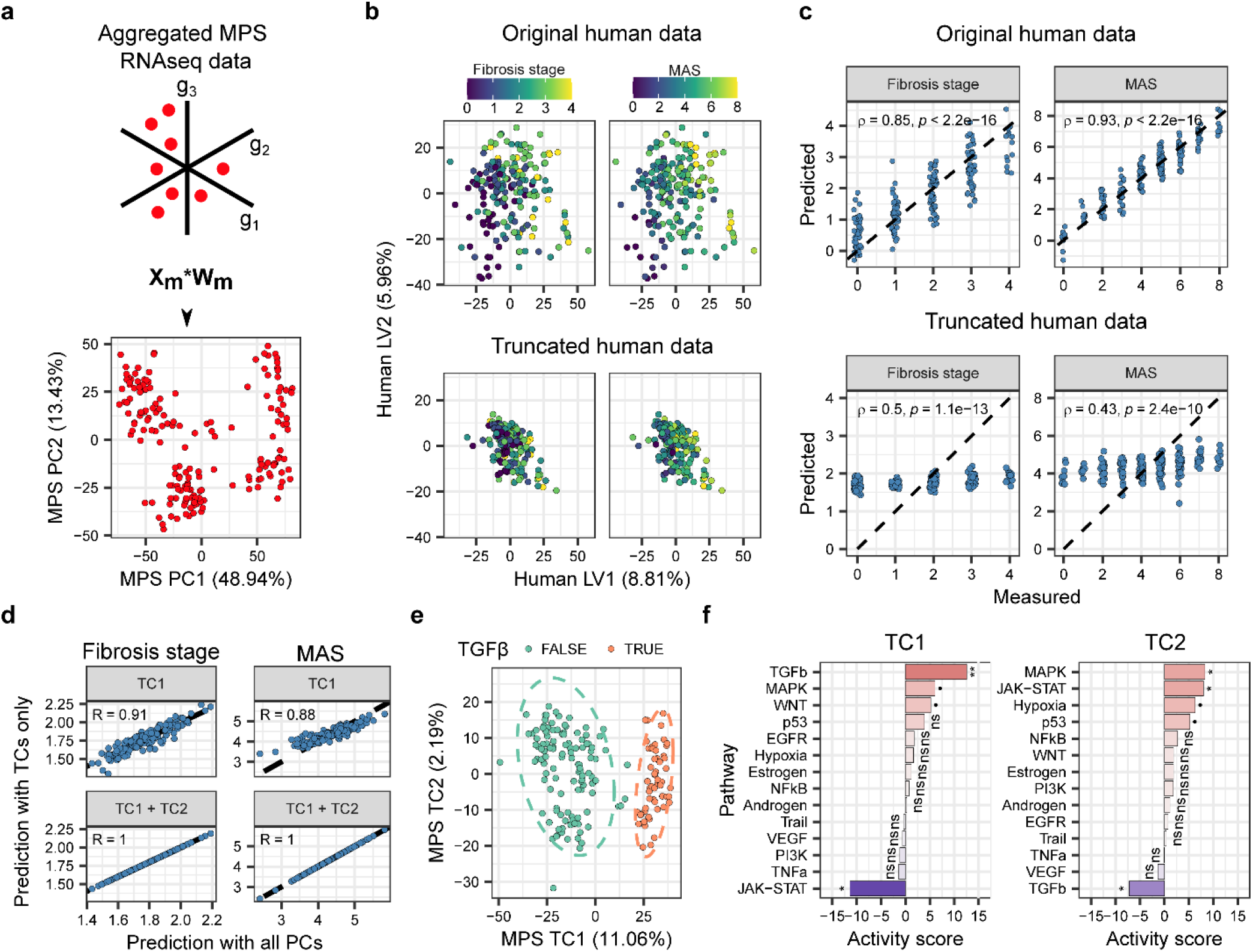
Translatable information in the MASH MPS is driven by TGFβ signaling. **(a)** Principal component analysis (PCA) is performed on bulk transcriptomics from the MPS model after aggregating experimental replicates. **(b)** Truncating human data through the MPS PC space highlights the loss of relevant biological variance. **(c)** The loss of biological variance leads to a decrease in the predictive power of the PLSR model. **(d)** Translatable information in the MPS PC space is contained in two translatable components that result in phenotype predictions identical to those made using the full PC space of the MPS model. **(e)** Projecting MPS data onto the two translatable components separates the samples based on whether TGFβ was included in the culture medium. **(f)** Estimation of pathway activity scores using gene loadings of the translatable components confirms that TGFβ signaling is significantly associated with the translatable components in the expected directions. Statistical significance for each pathway is calculated by counting how many times an equal high or higher absolute activity exists when using the PC loadings as input to the activity inference algorithm; asterisks indicate significance level defined as: ****p<=10−4, ***p<=10−3, **p<=10−2, *p<=0.05,·p<=0.1, and ns for p > 0.05.

### 2.4. Forward model translation highlights TGFβ responses as the main translatable factor in the liver MPS

While our approach reveals what predictive information is lost when truncating human data through the MPS-PC space, the resulting PLSR model still performs significantly better than a shuffled PLSR control (Supplementary Fig. 4), indicating that the MPS dataset still contains useful predictive information. We thus sought to find key directions in the MPS gene expression space that are responsible for preserving the translatable information. We defined translatable components (TC) as a set of orthogonal latent variables in the MPS gene expression space that can be used instead of the complete set of PCs in the data truncation step while achieving the same model predictions. TCs were calculated successively as linear combinations of MPS PCs to guarantee they contain non-zero MPS variance and that they are orthogonal to each other (Methods).

For the current case study, a set of two TCs was sufficient to achieve identical predictions to those obtained when using the complete set of MPS PCs (Figure 3d), with the first TC capturing most of the translatable information for both phenotypes. The two TCs accounted for around 13% of the total variance in the MPS dataset and did not correspond exactly to any of the MPS PCs (Supplementary Fig. 6), highlighting that the most translatable directions in the MPS dataset do not necessarily correspond to the directions of highest variance in the MPS data. Interestingly, projecting the MPS samples onto these two TCs resulted in two distinct clusters, which corresponded to conditions that included TGFβ or not, regardless of the other experimental stimuli (Figure 3e). Consistent with this observation, performing inference of pathway activity analysis on the gene loadings in the two TCs confirmed a strong TGFβ response signature in both TCs (Figure 3f). Taken together, these results suggest that for our chosen MPS, the addition of TGFβ to the culture medium was the added cue that resulted in most of the translatable information.

### 2.5. Reverse model translation reveals additional latent variables required to further tune MPS translation

After analyzing the source of translatable information in the MPS dataset, we next asked whether we could find unexplored directions in the MPS gene expression space that would improve information translation when truncating human data. We defined extra latent variables (LV Extra) as a set of latent variables orthogonal to the MPS PC space and to each other that represent directions where the MPS should have non-zero variance, but originally did not. Because PLSR models are linear, we were able to determine LV Extra analytically (Methods). For our case study, two additional latent variables were sufficient to retrieve the missing predictive information.

Projecting the human samples onto the two calculated LVs arranges the samples qualitatively based on their phenotypic scores, where most of the trend with the MASLD score was associated with LV Extra 1, while the fibrosis stage also varied along LV Extra 2 (Figure 4a). These observations are consistent with our optimization approach, where we first optimized for MASLD score with LV Extra 1 and then used LV Extra 2 to further tune for the fibrosis stage. Indeed, since fibrosis stage and MASLD are correlated (Figure 2b), LV Extra 1 was expected to also capture information relevant to estimating the fibrosis stage.

**Figure 4:**
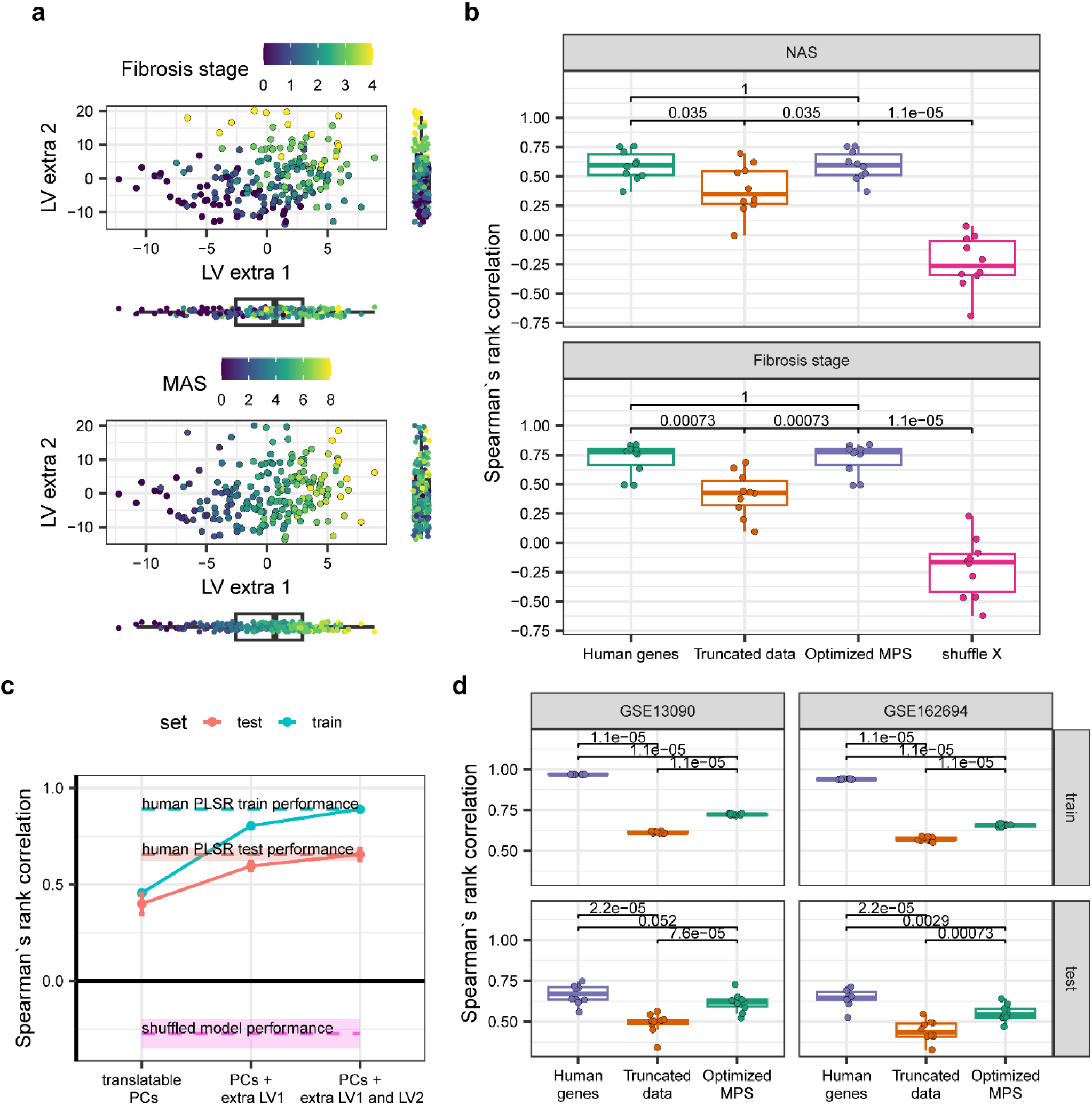
Additional latent variables not captured in MPS dataset are informative of MAFLD/MASH phenotypes. **(a)** Projecting human samples onto the set of two extra latent variables reveals qualitative trends in both phenotypic scores. **(b)** Appending the two extra latent variables to the PC space of the MPS significantly improves the prediction of both phenotypes compared to the PC space only (Truncated) and is equivalent to the performance of the human PLSR model. **(c)** Extra latent variables improve the prediction of phenotypes sequentially both in training and in testing, with the first extra LV accounting for most of the improvement. **(d)** Extra latent variables generalize to additional human datasets and improve the prediction of human phenotypes compared to using the PC space only (Truncated). For all comparisons in this plot, a two-sided unpaired Wilcoxon test was used. In all boxplots, the centerline denotes the median, the bounds of the box denote the 1st and 3rd quantiles, and the whiskers denote points not being further from the median than 1.5 × interquartile range (IQR).

Quantitatively, an augmented set of latent variables that included the full MPS-PCs and the two LV Extra (optimized MPS) performed significantly better on cross-validation than both a shuffled PLSR model and the PLSR model with truncation on just the MPS PC space, with no significant differences compared to the PLSR model with the full *in vivo* human dataset (Figure 4b). Because the LV Extra are calculated sequentially, they gradually increased the performance of the PLSR model in training and in testing until the performance of the full *in vivo* human dataset was achieved (Figure 4c). Importantly, the additional LVs also retrieved predictive information in PLSR models trained on independent human datasets (Figure 4d), suggesting that they are generalizable and represent disease biology not captured in the MPS dataset. The additional LVs found using PLSR are enough to retrieve the dropped performance of other machine learning models as well (Supplementary Fig. 5). Finally, we validated that for even slightly lower data partitions, we get similar extra latent variables and pathways loaded on them (Supplementary Fig. 7-8). Overall, this novel approach allowed us to identify potential directions in gene expression space where inducing variance in the MPS by applying new experimental cues can further tune the translatability of the MPS model.

### 2.6. Analysis of additional latent variables suggests a role for interferon response as an additional cue in liver MPS

While extra LVs are orthogonal to each other, they are calculated sequentially and are therefore not independent. In fact, any set of two orthogonal vectors spanned by the two extra LVs will be equivalent in terms of retrieving the missing human information. Therefore, we posed the question of whether a single extra LV corresponding to a linear combination of the two extra LVs calculated could capture most of the information that the two calculated LVs capture. We assessed the performance of the PLSR model when using a single additional LV defined as a weighted sum of the two calculated LV Extra (Figure 5a). As expected, using a mixed LV closer to LV Extra 2 decreased the correlation between the measured and calculated MASLD score since LV Extra 1 was defined to capture variance relevant to predicting the MASLD score. Interestingly, an intermediate mixed LV resulted in a higher correlation between predicted and measured fibrosis stages than that achieved with either LV Extra 1 or LV Extra 2 alone, which confirms that choosing a single extra LV leads to a trade-off between predicting either phenotype.

**Figure 5:**
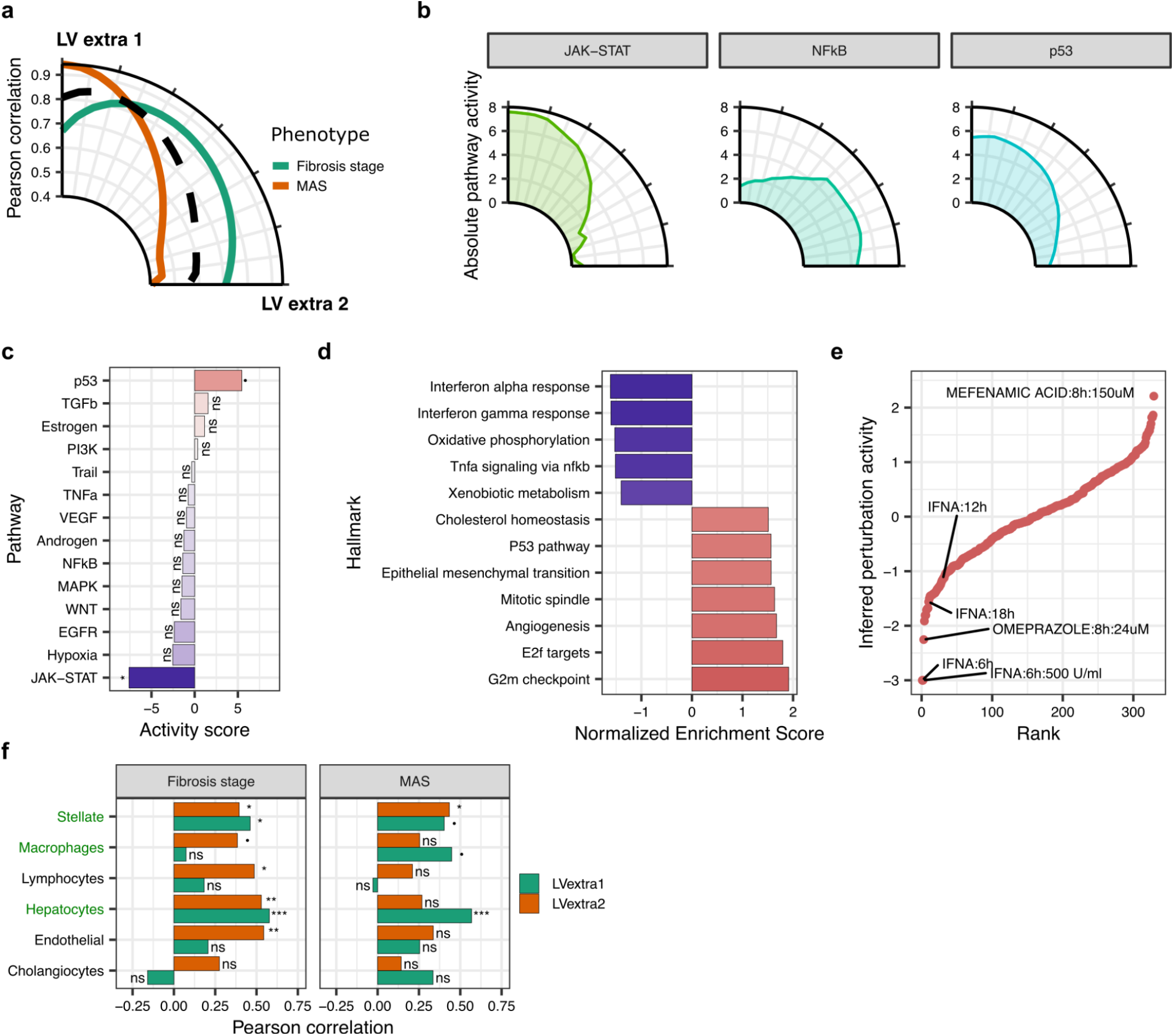
Analysis of extra latent variables highlights the role of interferons as a potential missing cue in the MPS model. **(a)** Selecting a single latent variable taken from the space spanned by the extra latent variables improves the prediction of human phenotypes on average, but there is a trade-off between predicting MAS and Fibrosis stage. **(b)** The region between the two extra latent variables that yields a good trade-off between the prediction of human phenotypes is characterized by high absolute activity of the JAK-STAT and p53 pathways and low activity of the NFkB pathway. **(c)** Inferred pathway activities that drive the observed transcriptomic profile are loaded in the first extra latent variable. Statistical significance for each pathway is calculated by counting how many times an equal high or higher absolute activity exists when using the PC loadings as input to the activity inference algorithm **(d)** Enriched Hallmark genesets loaded on the first extra latent variable. **(e)** Perturbations from ChemPert^33^ whose activity could induce the transcriptomic profile loaded in the first extra latent variable **(f)**. Human snRNAseq data pseudobulked by patient and major cell type was projected onto the two LV Extra, and the correlation between the resulting scores and the histological scores of the samples was calculated. Cell types annotated in green are present in the MPS used in this study. In all panels, asterisks indicate statistical significance level defined as: ****p<=10−4, ***p<=10−3, **p<=10−2, *p<=0.05,·p<=0.1, and ns for p > 0.05.

The calculated extra LVs revealed directions in gene expression space where increasing MPS variance can help preserve translatable information, but they do not indicate how much variance can be achieved. To aid with interpretation, we performed pathway activity inference^42^ on the gene loadings of the two extra LVs and on intermediate LVs (Figure 5b and Supplementary Fig. 9). This analysis revealed that the region in which a single mixed LV results in good model performance for both human phenotypes corresponds to directions in gene expression space dominated by the activity of the JAK-STAT and p53 pathways, while the contribution of the NFkB pathway is diminished. This result suggests that it may not be necessary to increase MPS variance exactly along the calculated LVs as long as the appropriate pathways, hence gene expression profiles, are perturbed.

Therefore, we focused on interpreting LV Extra 1, as it is the direction that provides most of the information retrieval for both the MAFLD score and fibrosis stage. Pathway inference analysis on this extra latent variable confirmed a significant association between its gene loadings and perturbation signatures along the p53 and JAK-STAT pathways with opposing signs (Figure 5c), indicating that any proposed experimental perturbation should perturb the JAK-STAT and p53 pathways in opposing directions. These results were corroborated by performing gene set enrichment analysis on the gene loadings, which highlighted a significant enrichment of genes associated with interferon response and cell cycle regulation along with other inflammatory and metabolic pathways (Figure 5d). To assess if LV Extra 1 was associated with known perturbagens, we compared its gene loadings with published transcriptional signatures curated from multiple non-cancer cell types^33^ (Figure 5e). Several perturbagens were identified with this approach, including interferons highlighted as agents capable of inducing gene expression signatures consistent with LV1 extra. Consistent with these findings, JAK/STAT signaling emerged as a relevant translational axis when repeating the analysis across additional in vitro-clinical dataset pairs (Supplementary Fig. 10).

Lastly, given that our reference MPS model only included hepatocytes, stellate cells, and macrophages, we verified that the lack of MPS variance along LV Extra could be due to the lack of a specific cell type present in human tissue. We used a human MAFLD/MASH single-nuclei RNAseq dataset^43^ and pseudobulked gene expression by patient for each annotated cell type (Methods). We then projected the resulting data onto the two calculated extra LVs to obtain projection scores and to calculate their correlation with the phenotypic scores for each patient. As anticipated, significant correlations between the projected scores and MAS were mostly limited to LV Extra 1, which was obtained by optimizing for MAS, while most of the significant correlations with LV Extra 2 scores were only apparent for the Fibrosis stage. Hepatocytes had the most significant correlations between both phenotypes and extra LVs, which is unsurprising because they comprise the bulk of the liver. Interestingly, endothelial cells, which are not present in the MPS model, displayed a significant correlation between fibrosis stage and their LV Extra 2 score, which suggests these cells might account for some of the variance along LV Extra 2 in human data that is not recapitulated *in vitro*.

Overall, these results suggest that including additional experimental perturbations, such as interferons, to increase MPS variance along a single additional LV can further tune the translatable information captured by the MPS model. Importantly, the lack of variance of the MPS data along LV Extras is not solely due to a lack of a specific cell type in the model, since hepatocytes, the most abundant cells *in vivo* and *in vitro*, contain important information along LV Extras in the human dataset analyzed. Therefore, additional cues can serve as a complement to other cues such as TGFβ, whose signaling response is already captured by the translatable components of the MPS (Figure 3f). The captured TGFβ signaling in the translatable components loadings, which was already identified as a key driver of the captured phenotypic variance in the MPS, serves as a validation step for the pathway inference approach and highlights that JAK-STAT signaling should be further investigated as an important pathway for *in vitro* to *in vivo* translation.

### 2.7. An unbiased translation approach identifies conditions to better capture human diversity in liver MPS

Beyond capturing disease-specific human biological variance, it might be desirable for *in vitro* models to also capture a broader range of human biological variance to model the potential adverse effects of new drugs. We developed an optimization approach (Figure 6a), utilizing stochastic gradient descent^44^, to maximize the total captured human variance by the MPS. Briefly (see Methods for more details), a differential gene expression vector (*ΔX*), which would be caused by a hypothetical experimental perturbation, is initialized randomly using a normal distribution with a mean and standard deviation equal to those of the MPS gene expression. This *ΔX* is then added to the centroid gene expression vector of the lean control samples in the MPS to yield a new gene expression vector (*X*^*^_*lean*_) that represents a new sample to be added to the data to maximize the human variance that can be captured by the MPS principal components space. Iteration proceeds, updating the differential gene expression vector (*ΔX*) until convergence, repeating the process with other random initializations and ultimately taking the average across all identified *ΔX* as the final determined perturbation. Adding just this one new sample results in increasing the human variance captured by the MPS PCs from ∼32% to ∼43%. To interpret the identified gene expression perturbations, we inferred the activity of pathways that can generate the calculated perturbation (Figure 6b). Interestingly, we observed that perturbing JAK-STAT signaling can also increase the total captured human variance, even though it is not the most prominent perturbation in this case. IFN-based responses are again found enriched when performing GSEA using hallmark genesets (Figure 6c), while transcription factors such as STAT2 and IRF9, downstream of JAK-STAT signaling, are highly ranked based on their inferred activity (Supplementary Fig. 11). Hypoxia-inducible factor 1-alpha (HIF1A) activity was also highly ranked (Supplementary Fig. 11), consistent with a significant enrichment of a response to hypoxia (Figure 6c). These suggest that inducing hypoxia in the *in vitro* model may prime the system to capture more of the human variance.

**Figure 6:**
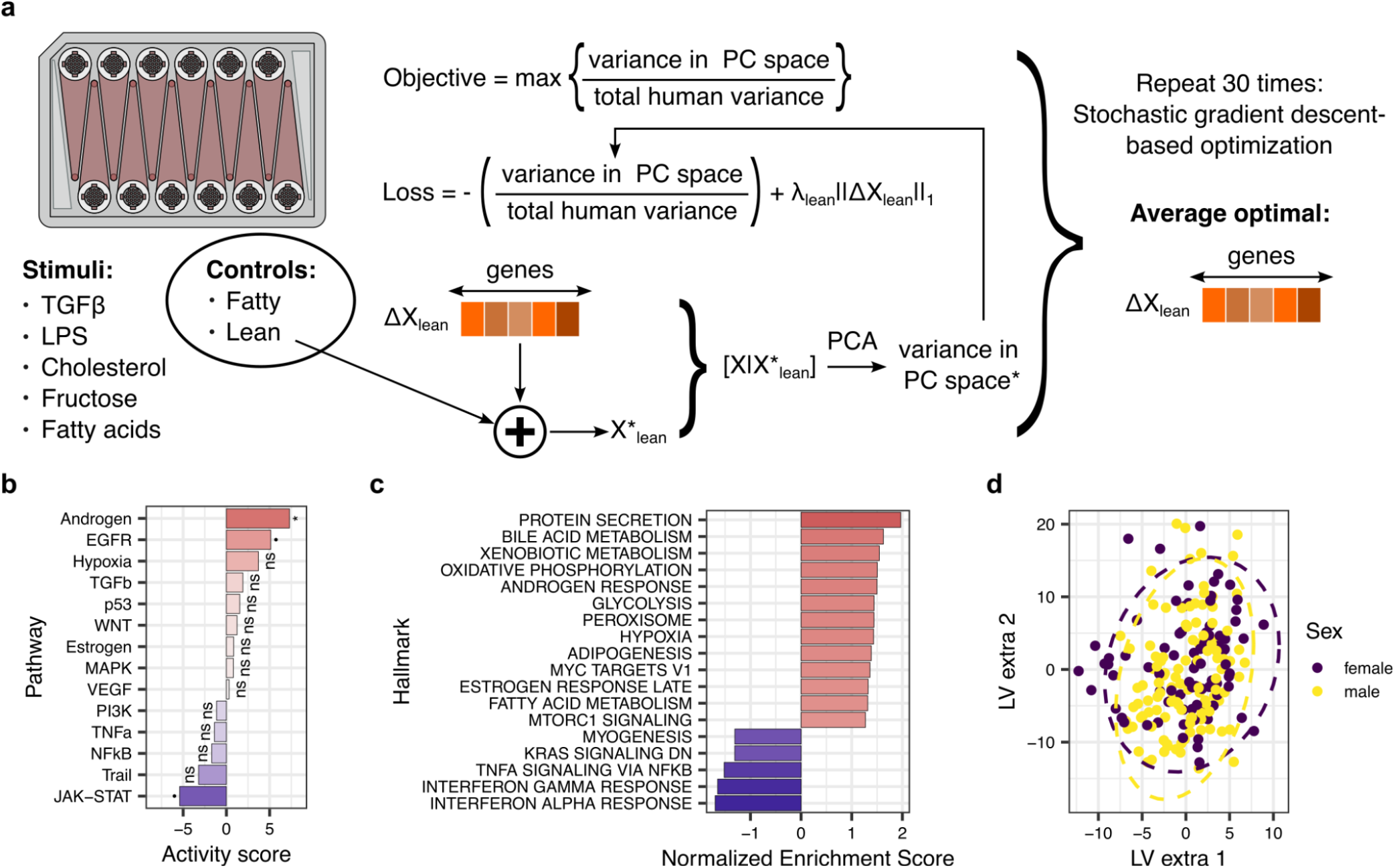
Identification of a differential gene expression vector for maximization of total MPS-captured human variance points to JAK-STAT signaling and sex-associated signals. **(a)** Schematic of the gradient descent-based algorithm for identifying a perturbation, in the form of a differential gene expression vector, that maximizes the total human variance captured by the MPS. **(b)** The induced pathway activity in the direction of the computationally derived perturbation points towards JAK-STAT, Androgen, and EGFR signaling. **(c)** The enrichment scores of the hallmark genesets loaded in the direction of the computationally derived perturbation. **(d)** Lack of sex separation of the samples in the identified extra latent space. Statistical significance for each pathway is calculated by counting how many times an equal high or higher absolute activity exists when using the PC loadings as input to the activity inference algorithm; asterisks indicate significance level defined as: ****p<=10−4, ***p<=10−3, **p<=10−2, *p<=0.05,·p<=0.1, and ns for p > 0.05.

Interestingly, the dominant effects to increase *in vitro* variance seem to be derived by perturbing primarily Androgen and secondly EGFR signaling (Figure 6b). Androgens, as well as estrogens, whose responses were enriched in the identified perturbation (Figure 6c), have been reported to be associated with both liver metabolism and MAFLD, as well as sex-associated differences in MAFLD, because of the difference in the activity and levels of androgens and estrogens between males and females^45–48^. Differences between male and female animals have also been observed regarding hepatic regeneration, dependent on androgen and estrogen signaling^45^, with the latter having a reported bidirectional relationship with EGFR^49^. The MPS in our case study likely contains only male samples (Supplementary Fig. 12-13), while the extra latent variables, which were calculated to better recapitulate MAS and fibrosis, do not explain the sex of the human patients (Figure 6d). Thus, we hypothesize that the identified perturbation, which maximizes total human variance, can potentially be equivalent to stimuli that can prime the *in vitro* system to be sensitive to sex-related differences. Alternatively, and perhaps more simply, additional MPS samples could be built using female-derived cells maintained in a female-specific culture medium.

## 3. Discussion

The primary objective of microphysiological systems (MPS) is to facilitate improved generation of preclinical information relevant for human contexts, reducing the dependency on animal models. In this study, we introduce a methodology that applies broadly across various *in vitro* models to identify translatable information within MPS. By leveraging transcriptomic data from both human and MPS systems, we employed a case study focused on Metabolic dysfunction-associated steatotic liver disease (MASLD). Our approach effectively retrieves the sources of key translatable information from a given MPS model and allows for prioritization of relevant experimental cues. Moreover, our method retrieves phenotypically relevant human variation by identifying extra latent variables pertinent to each phenotype of interest that allow for improved predictive capabilities of the MPS models, and suggests cues to achieve MPS variance along the identified latent variables. Importantly, our approach does not directly analyze the projection of human samples onto the MPS space as done in previous work^26,27^ and it goes beyond the idea of finding a gene expression subspace with good alignment between *in vitro* and human data^50,51^ because we also calculate important latent variables that are important for predicting human phenotypes regardless of whether they contain zero MPS variance. In this regard, our work represents a novel approach to systematically determining cues or perturbations that can be added to MPS models to improve their translatability without aiming to mimic human samples on a one-to-one basis.

For our particular case study, our results suggest that TGFβ serves as a crucial cue that preserves significant translatable information in the selected MPS. While the role of TGFβ in MASLD/MASH and fibrosis has long been recognized, our results highlight the potential of our method to prioritize experimental stimuli in other MPS models where key cues might not be as obvious. In contrast, the lack of additional information from other cues in the MPS may be attributed to the timing of sample collection. It is plausible that the MPS has adapted to initial perturbations by the time samples were collected for sequencing, therefore diminishing the net impact of specific cues. This observation aligns with the multi-hit theory of disease progression^52^, where diverse initial insults lead to a cascade of cellular events, fundamentally altering cell behavior and intercellular communication that ultimately dictate the phenotype of the tissue.

The identification of extra latent variables suggests that perturbations along the JAK-STAT signaling pathway – potentially mediated by interferon (IFN) – can further enhance the predictive accuracy of the MPS for human clinical phenotypes. The extra latent variables are calculated sequentially, and, in our case, we started by optimizing for the MASLD score, followed by tuning for the fibrosis stage. However, our overall conclusions are still valid if we choose to optimize first for the fibrosis stage and then tune for MASLD, since in this case, the resulting extra LVs are simply a rotation of the original extra LVs and therefore do not provide additional information. Notably, IFN-based perturbations of the JAK-STAT signaling pathway are identified by applying the approach also with other pairs of clinical-*in vitro* datasets, highlighting that this perturbation is potentially missing by many current *in vitro* models of the disease. IFNs have been shown to be involved in various inflammatory pathologies of the liver^53^. IFNγ is known to promote the inflammatory response of macrophages, which leads to the recruitment of additional immune cells, promoting inflammation and contributing to hepatocyte injury and fibrosis^39^. Meanwhile, type I IFNs are associated with the increased production of proinflammatory cytokines in the liver. Thus, the involvement of IFNs in the disease development and progression not only makes them interesting potential therapeutic targets but also potentially useful stimuli to be added to the media conditions of the *in vitro* model, to better mimic the tissue microenvironment of MAFLD. However, proper experimental validation of these ideas is necessary to assess their value.

In the context of recapitulating overall human variance *in vitro*, our numerical algorithm revealed once again a potential role for JAK-STAT signaling. These results are consistent with the calculated extra LVs and are not surprising given that this particular human dataset was collected with the purpose of studying MASLD/MASH, and we therefore expect that a significant proportion of human variance in the data is directly associated with disease biology. However, our results highlighted a lack of potentially sex-related transcriptional responses in the MPS model, which was made with mostly male-derived cells. Given that sex differences in MAFLD have been described^45–48^, our results strengthen the need to include sex as an important variable in MPS studies^54^.

A limitation of our approach is that it requires molecular features measured across a wide range of experimental conditions to produce a good assessment of how much disease-relevant biological variance a particular MPS can capture. This is because our method relies on leveraging covariance structures in molecular data. However, collecting molecular data from at least two conditions (*e*.*g*., control vs disease) is enough in principle to calculate a single latent variable that can then be used to calculate additional latent variables required to capture more of the human-relevant biological variance. Our approach can be integrated as part of an iterative cycle of perturbing MPS models and assessing translatable information to refine *in vitro* models progressively. Notably, we relied solely on gene expression data, mainly due to data availability and their high predictive power for the case study of MAFLD. It is possible that other types of molecular features (i.e. proteomic, metabolomic) could provide complementary information that would lead to different assessments of MPS quality. Moreover, the predictive ML model used can be extended to a multi-omic approach that simultaneously accounts for different levels of molecular profile and their interactions, inspired by the Latent Interacting Variable Effects (LIVE) modeling^55^.

Overall, we anticipate our approach, LIV2TRANS, will be useful in addressing the question of how complex an MPS model needs to be to model a disease of interest accurately. It provides a principled way for selecting cue conditions that align with relevant phenotypic directions in humans, thereby advancing our understanding of disease mechanisms and their therapeutic solutions, and accelerating the process of *in vitro* model design. Our method builds on the premise that, instead of attempting to design complex *in vitro* models that allow for simple one-to-one mapping to human samples, *in vivo* response can be built as a function of responses in diverse *in vitro* models that are, on the whole, informative because they capture important biological variance^56^.

## 4. Materials and methods

### Datasets

Publicly available human MAFLD liver and MPS transcriptomic datasets were used in this study. The main clinical dataset used for the development of our pipeline (Govaere et al^30^) can be obtained from the Gene Expression Omnibus (GEO) with the number GSE135251. Two additional datasets used for model validation (Hoang et al^32^, Pantano et al^31^) can be obtained from GEO with accession numbers GSE13090 and GSE162694, respectively. Sample metadata was directly obtained from GEO. However, to reduce data variability due to different preprocessing bioinformatic pipelines, clinical datasets were downloaded as gene counts from ARCHS4^57^, which guarantees uniform preprocessing from raw sequencing files. The liver MPS dataset^8^ was not available in ARCHS4 and was obtained as gene counts along with sample metadata from GEO with accession GSE168285. Single nuclei RNA sequencing from human livers across MAFLD stages of disease^43^ were downloaded as an annotated Seurat file with accession number GSE202379.

### Model translation pipeline

#### Data preprocessing

For a given pair of human/MPS datasets, only common features (i.e. genes) found in both datasets were included. Gene counts were then normalized for each sample in each dataset as counts per million (CPM) and normalized as log2(1 + CPM).

#### Partial least squares modeling of human MAFLD/MASH

Partial least squares regression (PLSR) was used to build a model to jointly predict MAFLD scores and Fibrosis stage in the human dataset as a function of normalized gene expression data. Eight latent variables were determined to be optimal based on cross-validation. The PLSR model can be represented as *Y*_*h*_ = *X*_*h*_*W*_*h*_*B*_*h*_, where *Y*_*h*_ is a centered matrix of human phenotypes (samples by phenotypes), *X*_*h*_ is a matrix of normalized and centered human gene expression data (samples by genes), *W*_*h*_ is a matrix of weights (genes by latent variables) and *B*_*h*_ is a matrix of regression coefficients (latent variables by phenotypes). The obtained PLSR model was initially tuned using 10-fold cross-validation (using 80% of the data from Govaere et al^30^) and evaluated in a smaller test set (20% of the data), for potential over-fitting in the validation set, due to the selection of a specific number of the latent variables. Afterward, the number of latent variables was set to 8, and for the rest of the comparisons, the obtained model was trained and evaluated using a new 10-fold cross-test split of all the data from Govaere et al^30^. One fold (10%) of the data was hidden during training and used to evaluate performance in unseen data, while 90% of the data was used for training.

#### Translation of human data through MPS data

Principal component analysis (PCA) was applied to the normalized and centered MPS gene expression data (*X*_*m*_) after averaging the replicates for each experimental condition based on sample metadata. The averaging of replicates was performed so that the PCA model only captured variance in the data associated with the experimental conditions, and not with technical variability due to experimental or sequencing variability. The rotation matrix of the PCA model (*W*_*m*_) contains the principal components of the data, which are a set of orthonormal vectors that span the subspace in gene expression space that captures the full variance of the MPS dataset.

Human gene expression was then projected and back-projected onto the MPS PCA space as 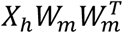, where the superscript *T* denotes the transpose of the matrix. The first projection onto the MPS space effectively truncates components of the human gene expression data that are not captured by the MPS dataset, while the back-projection places the data again in human gene expression space, which is necessary so that this truncated human data can be used with the human PLSR model. This truncated, or filtered, data is then evaluated with the trained human PLSR model to obtain a prediction of human phenotypes as 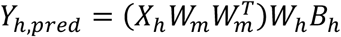.

#### Calculation of translatable components

Translatable components were defined as latent variables in the MPS space that preserved most of the translatable information contained in the full rotation matrix *W*_*m*_. That is, we are looking for vectors *w*_*TC*_ that result in similar predictions to those obtained with the complete set of MPS PCs and therefore 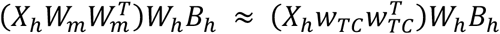. To guarantee that *w*_*TC*_ contain non-zero variance of the MPS data, translatable components were defined as a linear combination of MPS PCs as *w*_*TC*_ = *W*_*m*_*α*_*TC*_ with weights *α*_*TC*_. The problem now is to find a suitable *α*_*TC*_ such that 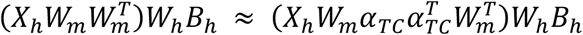. Defining 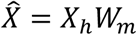 and 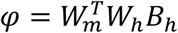 we transform the previous expression into 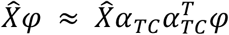. For a single phenotype, hence single *B*_*h*_ and *φ*, the previous expression becomes an exact equality if *α*_*TC*_ = *φ*/‖*φ*‖_2_, with ‖*φ*‖_2_ denoting the Euclidean norm of *φ*.

When more than one phenotype is present, each phenotype has a *α*_*TC*_ associated with its particular *B*_*h*_. In this case, any set of orthonormal vectors spanned by *W*_*TC*_ = *W*_*m*_*A*_*TC*_, where *A*_*TC*_ is a matrix containing the *α*_*TC*_ of each phenotype as columns, is a valid space of translatable components. Therefore, to aid with interpretation and to guarantee that the order of translatable components is meaningful, we performed a line search to get an intermediate vector *w*_*TC*_ between the two vectors *W*_*m*_*A*_*TC*_ that minimized an overall normalized prediction error 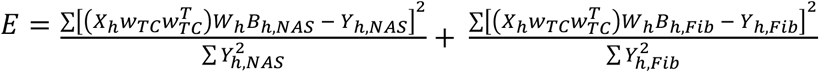. The intermediate vector with the lowest prediction error was defined as the first translatable component *w*_*TC*,1_. The second translatable component *w*_*TC*,2_ was then calculated as a vector orthogonal to *w*_*TC*,1_ such that both translatable components span the same space as *W*_*TC*_ = *W*_*m*_*A*_*TC*_. This approach can be extended to cases with more than two phenotypes by performing successive rounds of optimization of a prediction error, although direct line searches will become more inefficient in this case.

#### Calculation of additional latent variables

Additional, or extra, latent variables were defined as latent vectors orthogonal to the MPS space that contain translatable human information. These variables were calculated analytically by noting that the PLSR model is a linear model with *Y*_*h*_ = *X*_*h*_*W*_*h*_*B*_*h*_ = *X*_*h*_*φ*_*PLS*_, where *φ*_*PLS*_ = *W*_*h*_*B*_*h*_ is a matrix of size genes by number of phenotypes. Therefore, the phenotypes predicted when including the projection-backprojection step are 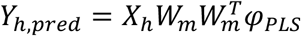. Extra latent variables *w*_*extra*_ are set of vectors that when appended to *W*_*m*_ form a matrix 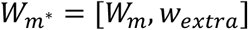 such that 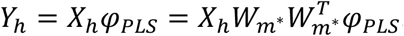. The vectors *w*_*extra*_ are defined to be unit-norm and orthogonal to each other and to *W*_*m*_ such that they provide information not contained in the MPS, which implies that matrix 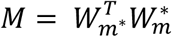 equals an identity matrix and that any linear combination *λ* of the columns of 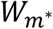 results in 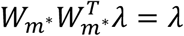. Therefore, for 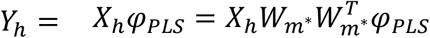 to hold, *φ*_*PLS*_ must be in the column space of 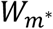. To solve this problem, *φ*_*PLS*_ was regressed against the MPS PCs *W*_*m*_ such that *φ*_*PLS*_ = *W*_*m*_*β* + *ε*, where *ε* is a residual term. Solving the least-squares problem results in 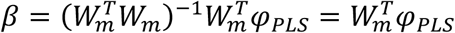 and 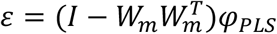 where *ε* is orthogonal to all columns of *W*_*m*_. Therefore, *w*_*extra*_ = *ε*/‖*ε*‖_2_ since *φ*_*PLS*_ = *W*_*m*_*β* + *w*_*extra*_‖*ε*‖_2_ is exactly a linear combination of the columns of 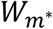.

Since each phenotype in the model has a vector *B*_*h*_ associated with a distinct *φ*_*PLS*_, any set of vectors that span the space spanned by the *w*_*extra*_ calculated for each phenotype is a valid set of LV Extra, although in general these *w*_*extra*_ will not be orthogonal to each other. To solve this issue, another possibility is to calculate *w*_*extra*_ for one phenotype and define this as LV Extra 1. Then augment *W*_*m*_ with this vector and calculate *w*_*extra*_ again with the *φ*_*PLS*_ of a second phenotype using the augmented *W*_*m*_ which guarantees that the new *w*_*extra*_ is orthogonal to *W*_*m*_ and to LV Extra 1, and so forth until all phenotypes have been examined. This latter procedure was used in the MASL/MASH case study since both the MAS score and Fibrosis score are correlated, and therefore LV Extra 1 calculated from the MAS score also retrieves information relevant to Fibrosis, while LV Extra 2 retrieves the remaining information.

### Unsupervised optimization of translatable variance

To identify a perturbation to maximize the human gene expression variance that is captured by the MPS model, a differential gene expression vector, that is added to a sample of interest is tuned using stochastic gradient descent^44^. Specifically, the mean of the lean control samples in the MPS is selected as a starting point, and a vector (*ΔX*), with dimension the same as the number of genes, is randomly initialized from a standard normal distribution. The Adam optimizer^58^ in the PyTorch framework^59^ (version 1.12) is picked to perform optimization, using a learning rate of 0.1 and 1000 iterations for the optimization process. During the optimization process, the goal is to maximize the variance that is captured by the MPS principal components space, through the minimization of the objective function: 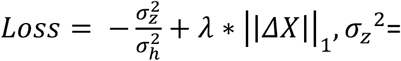 variance of the projected human samples, *σ*_*h*_^2^= total variance of human gene expression. The second term applies L1 regularization on the learnable differential gene expression vector (*ΔX*). The process is repeated 30 times to account for the random initialization, and the average of all the learned vectors is taken as the identified differential gene expression perturbation that can maximize the captured human variance.

### Pathway and signature analysis of latent vectors

The activity of pathways and transcription factors (TFs) that can drive the observed gene expression or contributions, was inferred from transcriptomics data using the VIPER algorithm^60^ from the decoupleR resource^42^. The VIPER algorithm calculates the enrichment of known source entities (such as TFs or pathways), which act as proxies of TF and pathway activity. The activity of a TF is calculated based on the expression of downstream genes known to be regulated by this specific TF, utilizing a known transcription regulatory network, while for the pathways the effect after perturbation on known downstream genes is used as a regulon. Specifically, for the TFs the Dorothea regulon^61^ was utilized with interactions of confidence levels A and B. For the pathways, the network describing the effect of 14 pathways to downstream genes, which is contained in the PROGENy resource^62^ was used.

### Gene Set Enrichment Analysis (GSEA)

Gene Set Enrichment Analysis (GSEA) was performed using the FGSEA library^63,64^ from the Bioconductor resource^63^. Thus, the genes’ contributions to the identified vector basis, differential gene expression vectors, as well as the gene-level profile of each perturbation, were transformed into a gene set-level profile of Normalized Enrichment Scores (NES). Statistical significance is calculated internally in FGSEA by calculating how likely it is to observe an equally high or higher enrichment score randomly based on a null distribution of enrichment scores, constructed by randomly permutating the data 10000 times. FGSEA is also performing p-value adjustment using the Benjamini-Hochberg (BH) correction.

### ChemPert-based perturbation inference

To identify perturbations that can drive the observed gene contribution profiles of the principal component space, the translatable components, the extra basis, and the differential gene expression vector that maximizes the captured human variance, the VIPER algorithm^60^ from the decoupleR resource^42^ was used coupled with a drug to transcription factors (TFs) regulatory network, contained in the ChemPert database^33^. The TFs activity is first inferred as already described and is used as an observed downstream signal that is regulated by the perturbations. The absolute value of the inferred perturbation activity is used as a proxy for the prioritization of conditions for subsequent experiments.

## Data and Code availability

The study did not produce new experimental data. All preprocessed data that were used to train our models, construct the ML framework and produce all tables and figures, as well as all the code to generate those data, figures and tables are available at the following GitHub repository: https://github.com/NickMeim/FattyLiverModeling.

## Acknowledgments

The authors would like to thank Brian Joughin for his valuable and rigorous input on this work. We also would like to thank Hratch Baghdassarian, Diana Gong, Christine Wiggins, Krista Pullen, and Luka Karginov for their valuable feedback. We acknowledge funding from the US Army Research Office Cooperative Agreement W911NF-19-2-0026 (DAL).

## Author contributions

JLC, NM, and DAL conceived the study. JLC preprocessed the data. NM implemented and executed cross-validation analyses of machine learning models. JLC and NM developed the mathematical and computational framework to identify cues that enhance *in vitro* models. JLC and NM wrote the manuscript and generated the figures. TM and ET performed experimental validation, generated figures for the experimental results, and edited the manuscript. DAL and LGG edited the manuscript.

