## Supplementary Material for "Systems biology framework for rational design of operational conditions for *in vitro / in vivo* translation of microphysiological systems"

* These authors contributed equally.

### Supplementary Figures

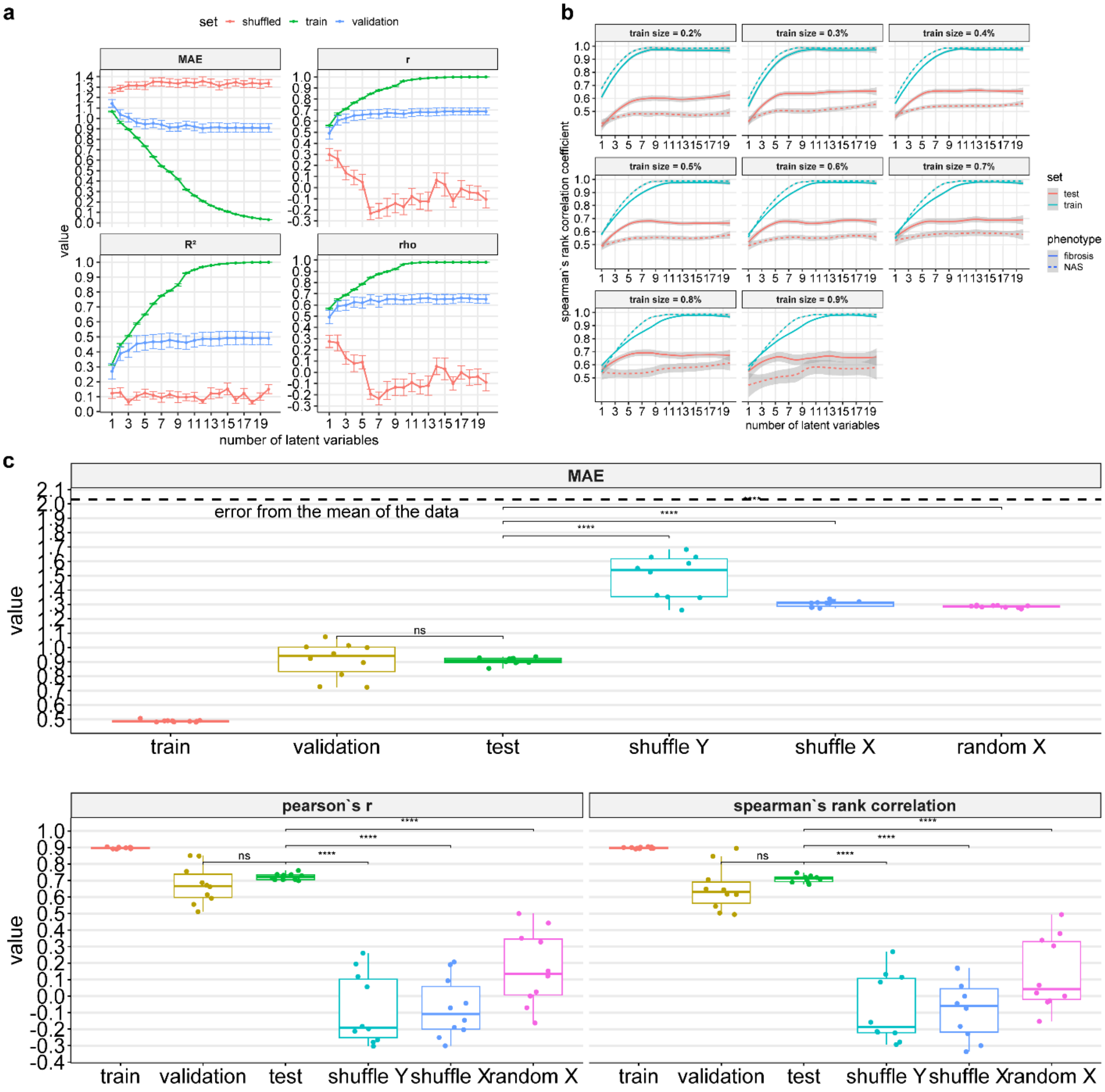


**Figure S1: Selecting the number of the latent variables in PLSR using 10-fold cross-validation on 80% of the data, while keeping 20% as a test set. a)** The effect of the number of latent variables in the performance of the PLSR model in the training and validation sets, in terms of mean absolute error (MAE), Pearson's correlation (r), coefficient of determination ($R^2$) , and Spearman's rank correlation coefficient (rho). The error bars denote a deviation of one standard error (SE) from the mean. **b)** The effect of the percentage of data used for training, in combination with the number of latent variables, on the performance of the PLSR model. The shaded area represents a deviation of one SE from the mean. **c)** Comparison of the performance of the PLSR model in training, validation, test set, as well as in the test set when using three randomized models, derived by shuffling the outputs (Y), the inputs (X), and using random features (X) during training. In all boxplots, the centerline denotes the median, the bounds of the box denote the 1st and 3rd quantiles, and the whiskers denote points not being further from the median than 1.5 × interquartile range (IQR).

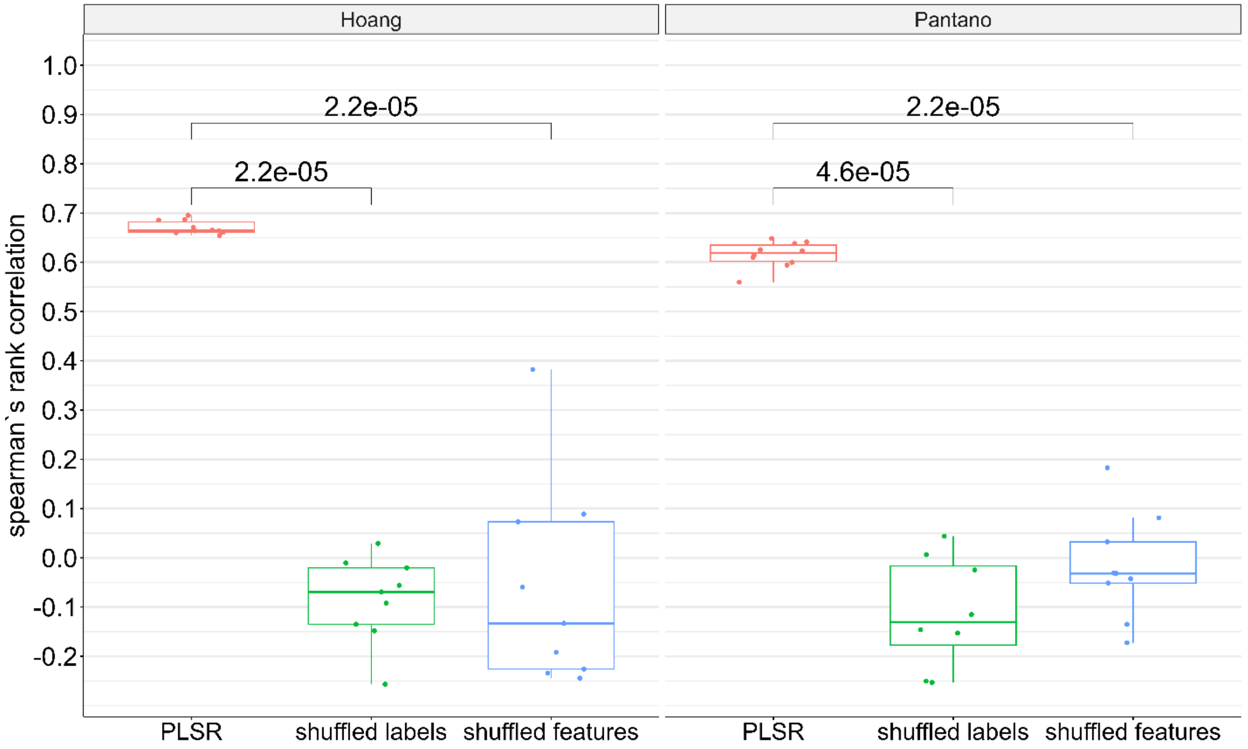


**Figure S2: Performance of the PLSR model (with 8 latent variables) in two external clinical datasets.** Each clinical dataset [1,2] is used as an external test set and each of the trained PLSR models, in each of the folds of the 10-fold cross-test approach, is used for making predictions. Performance is compared with two randomized models derived by shuffling the labels of the phenotypes, and the input features during training. In all boxplots, the centerline denotes the median, the bounds of the box denote the 1st and 3rd quantiles, and the whiskers denote points not being further from the median than 1.5 × interquartile range (IQR). The statistical test used to estimate whether the performance changes, within each data partition, is a two-sided paired Wilcoxon test.

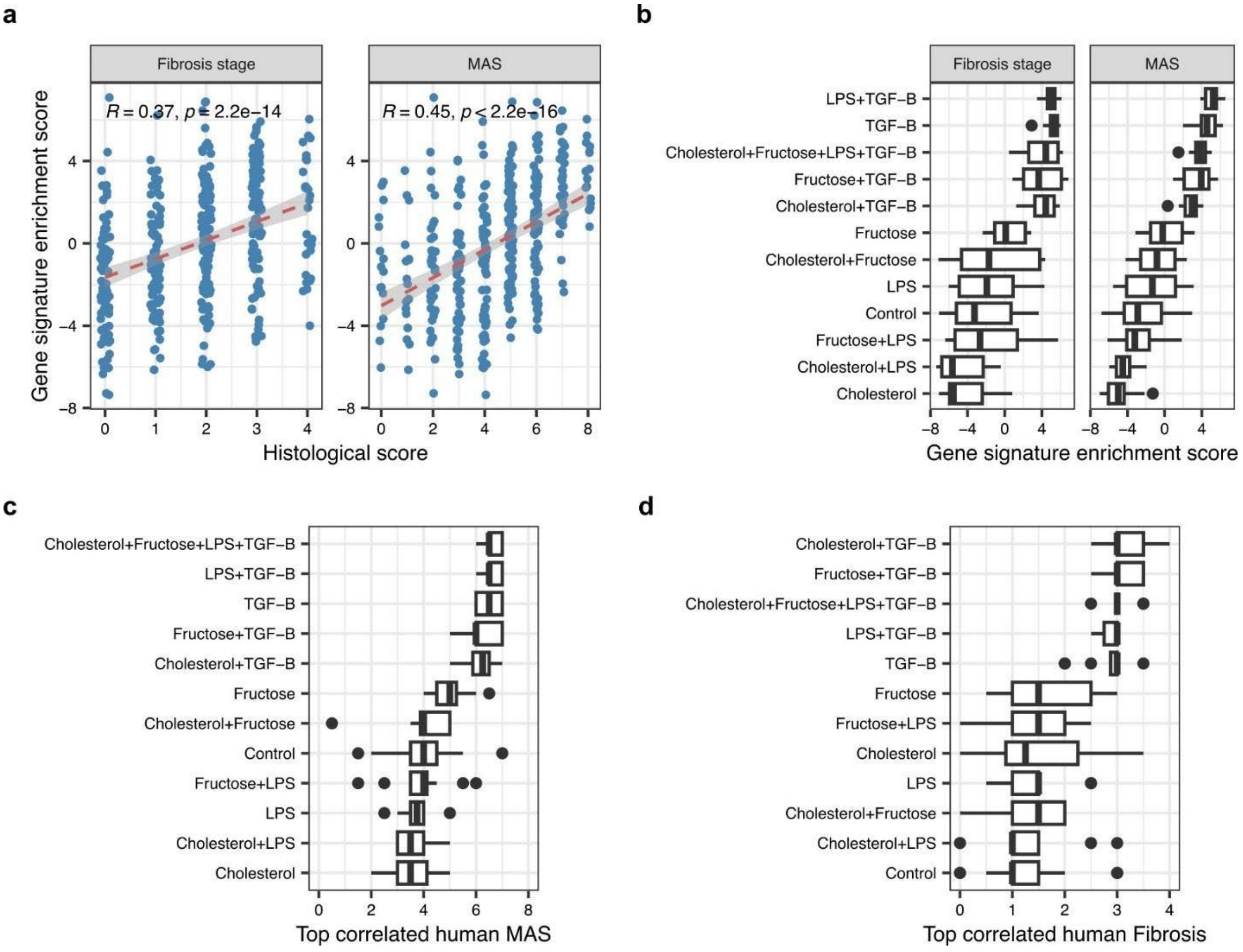


**Figure S3: Gene signature enrichment and sample-to-sample correlations between MPS and human data.** Enrichment of a published and validated gene signature for MAS and Fibrosis [1] was calculated for human RNAseq data (a) and for MPS data (b) using the non-parametric VIPER algorithm [3]. Sample-to-sample correlation was calculated between the normalized and centered human and MPS samples. For each MPS sample, as defined by their experimental design, the MAS score (c) and Fibrosis stage (d) of the top 10 highly correlated human samples are plotted.

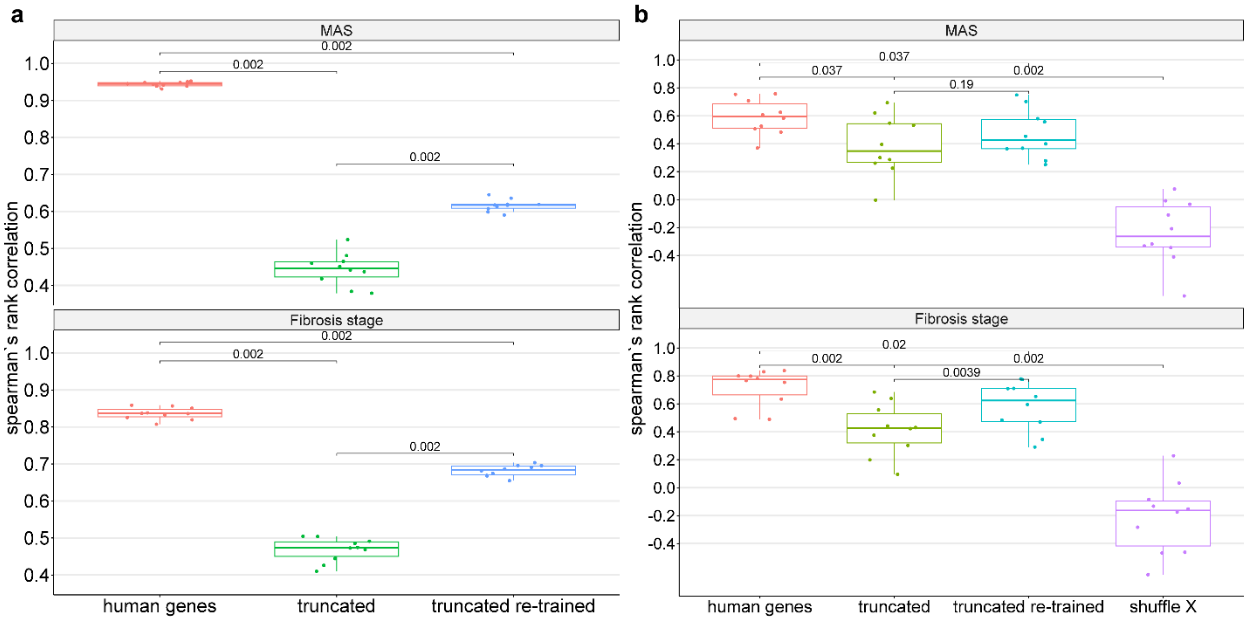


**Figure S4: Comparison of the performance of the PLSR model when using the original human genes as input, the genes truncated through the MPS space, and the truncated genes with the coefficients of the regression re-trained. a)** Performance in training when using a 10-fold cross-test approach. **b)** Performance in the test set of the 10-fold cross-test approach, while also comparing with a randomized model trained on shuffled features (X). In all boxplots, the centerline denotes the median, the bounds of the box denote the 1st and 3rd quantiles, and the whiskers denote points not being further from the median than 1.5 × interquartile range (IQR).

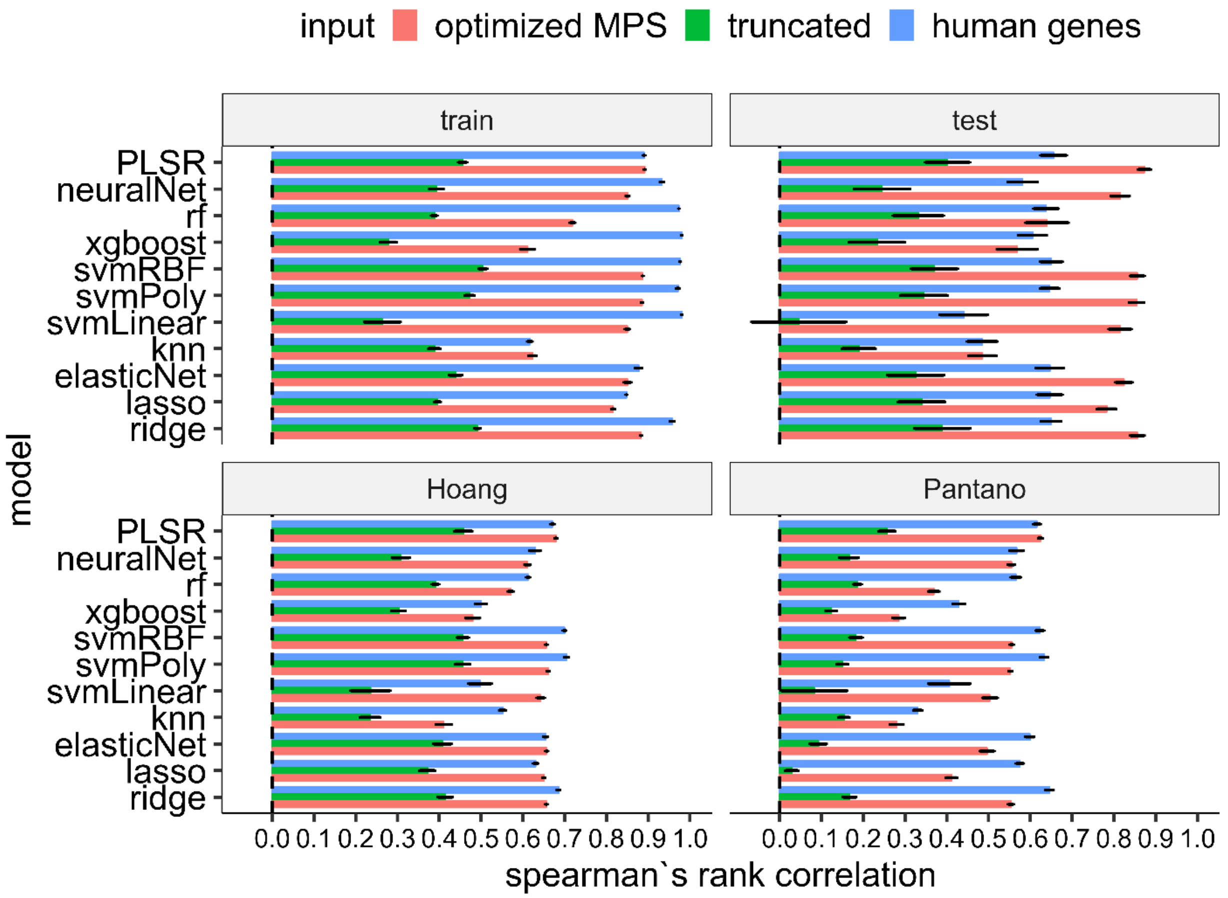


**Figure S5: Comparison of the performance of different machine learning models when using the original human genes as input, the genes truncated through the MPS space, and the truncated genes through the optimized MPS space.** The comparison is done by using a 10-fold cross-test and further validating models' performance on true external clinical datasets. For the optimized MPS space we utilized the same extra basis vector that was found using all the data in PLSR. The error bars denote a deviation of one standard error (SE) from the mean.

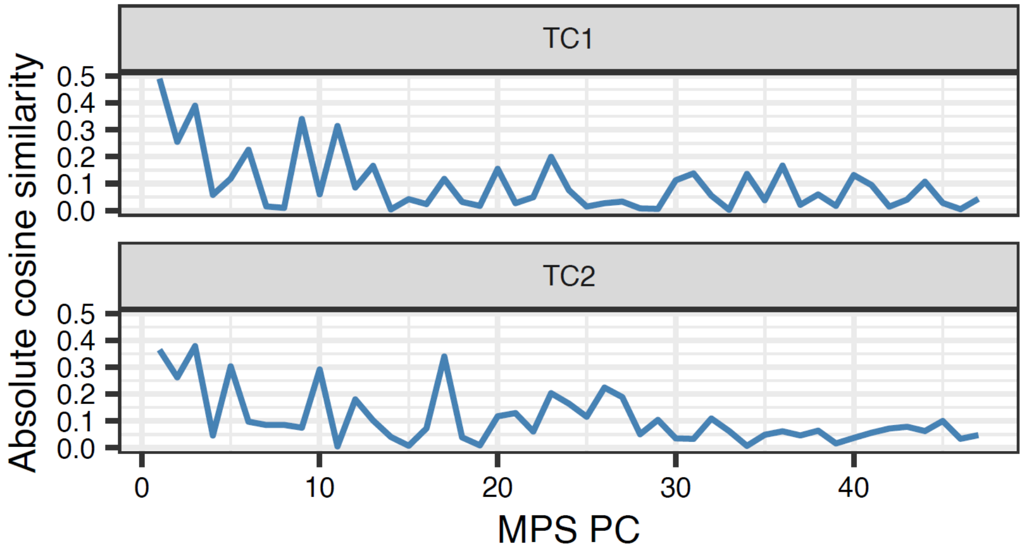


**Figure S6: Absolute cosine similarity between each translatable component (TC) and each principal component (PC) of the MPS dataset.** Absolute cosine similarity less than one shows that none of the PCs of the MPS dataset matches either of the TCs exactly.

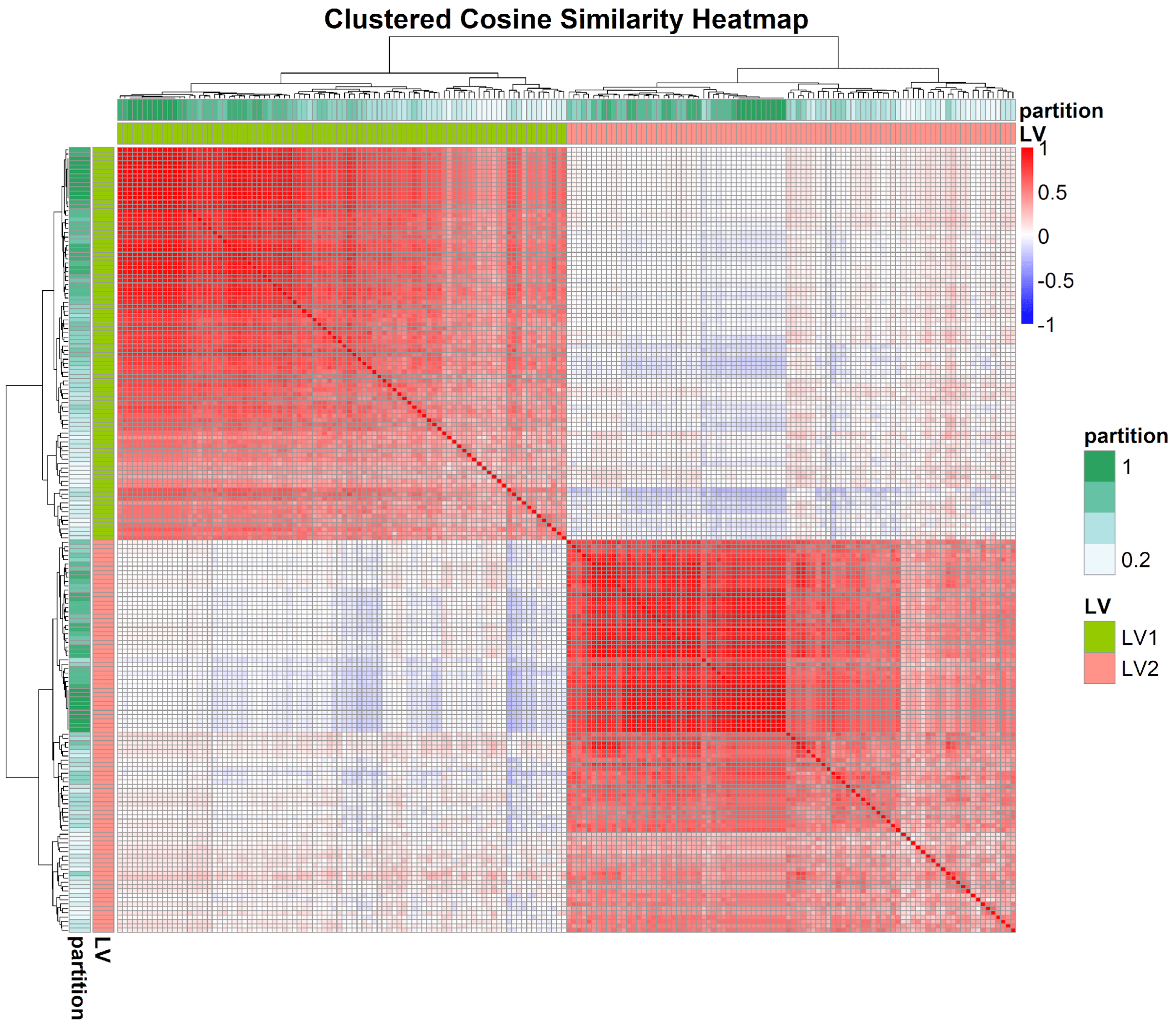


**Figure S7: Clustermap based on cosine similarity of gene loadings on extra LVs for different data partitions.**

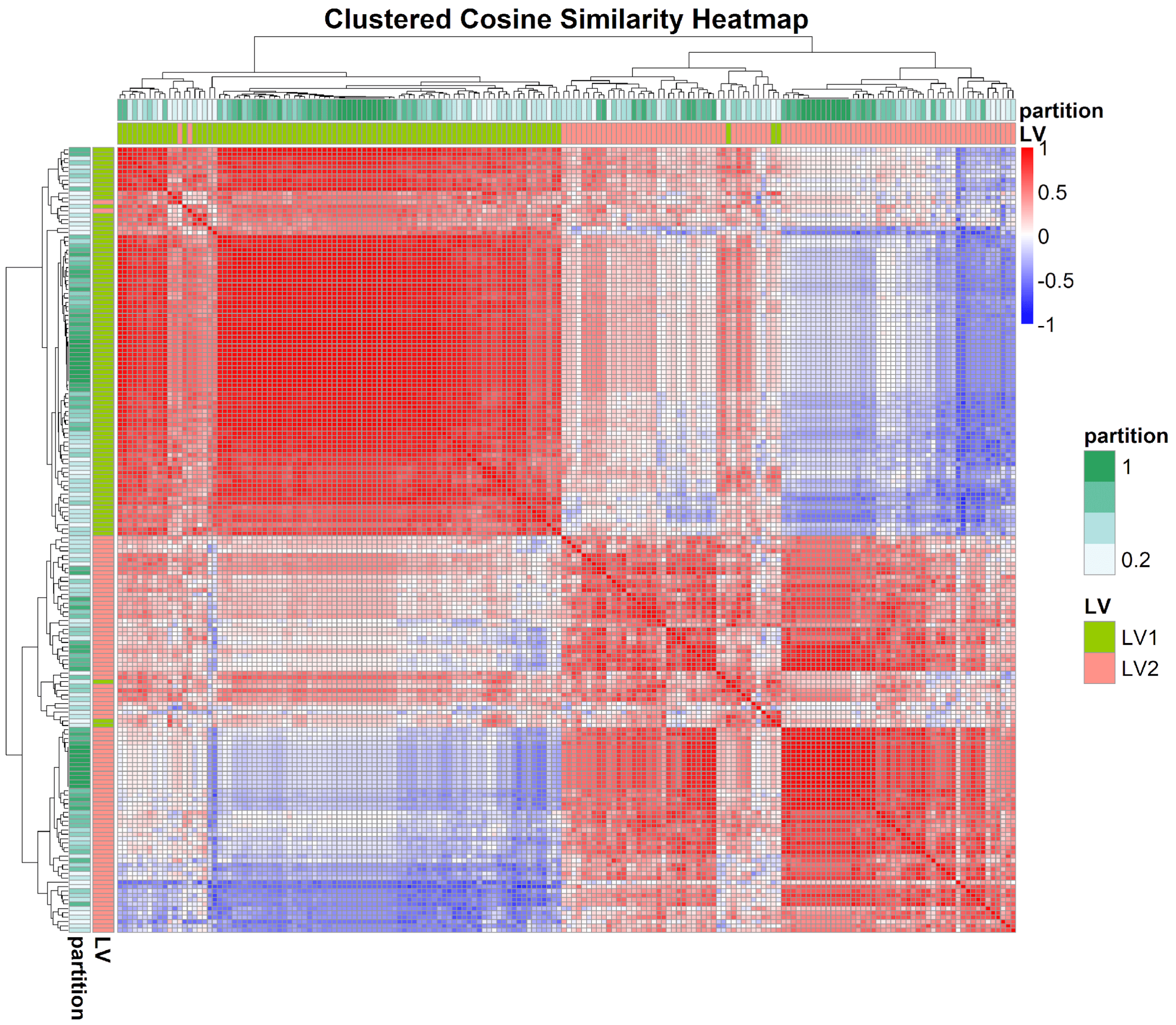


**Figure S8: Clustermap based on cosine similarity of pathway loadings on extra LVs for different data partitions.**

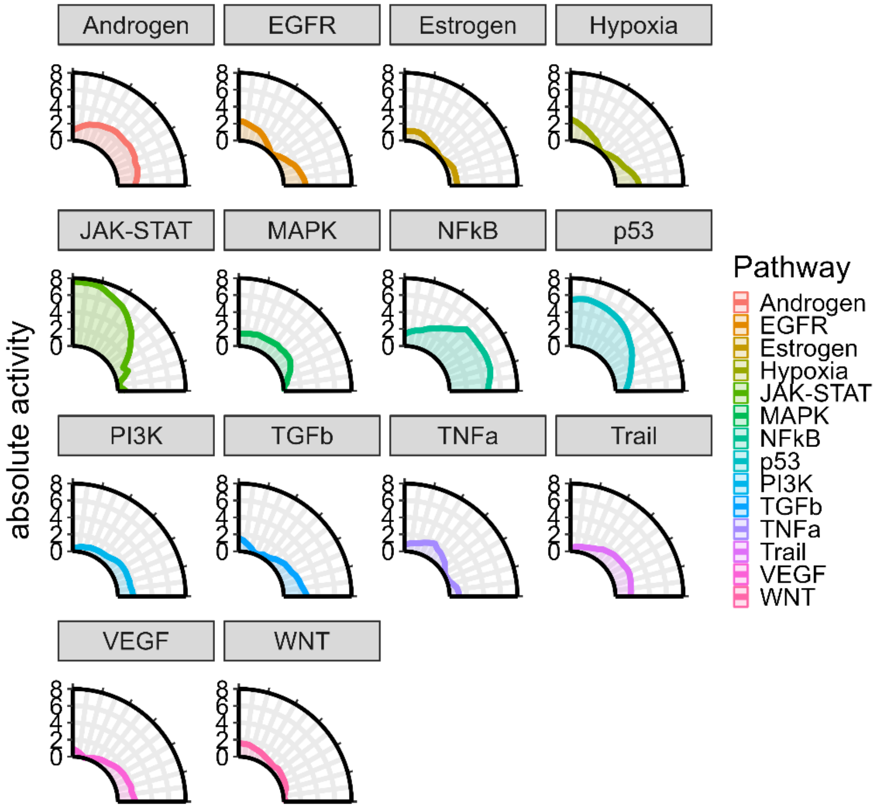


**Figure S9: Absolute activity for different linear mixing between extra LV1 and extra LV2.** The region between the two extra latent variables that yields a good trade-off between the prediction of human phenotypes is characterized by high absolute activity of the JAK-STAT and p53 pathways and low activity of the NFkB pathway, while the rest of the pathways do not seem to contribute significantly for any combinations.

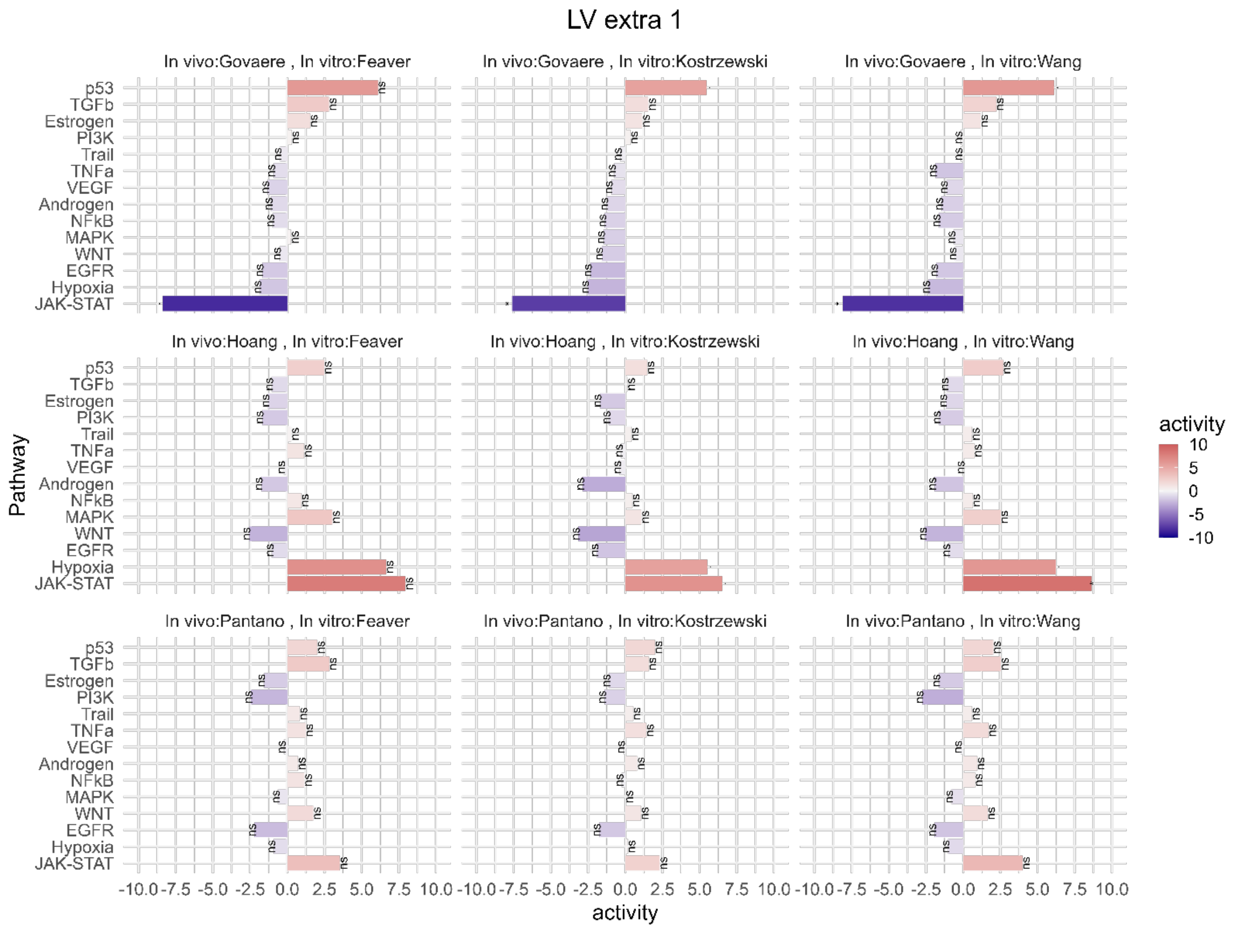


**Figure S10: Inferred pathway activities that drive the observed transcriptomic profile loaded in the first extra latent variable, using different pairs of clinical and in vitro datasets.** Statistical significance for each pathway is calculated by counting how many times an equal high or higher absolute activity exists when using the PC loadings as input to the activity inference algorithm

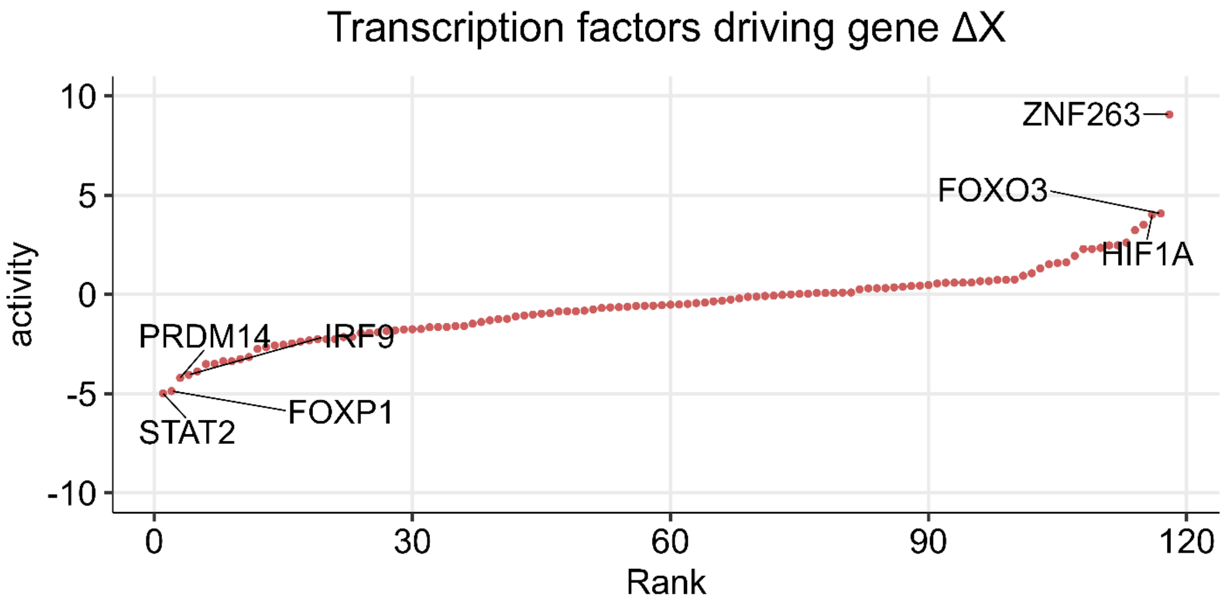


**Figure S11: Ranked TFs that are highly loaded in the perturbation that maximizes the total human variance captured by the MPS.**

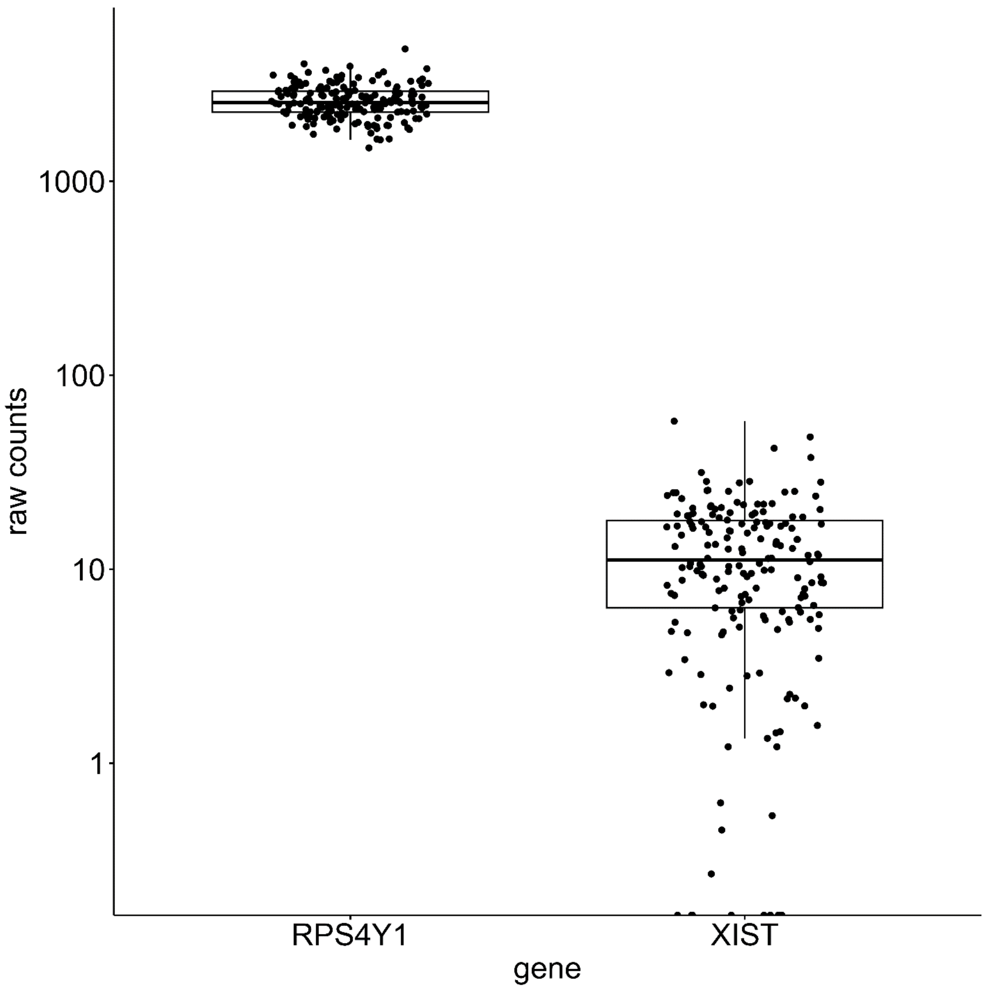


**Figure S12: Raw counts of genes that can be used to differentiate between males (RPS4Y1) and females (XIST) in the MPS.**

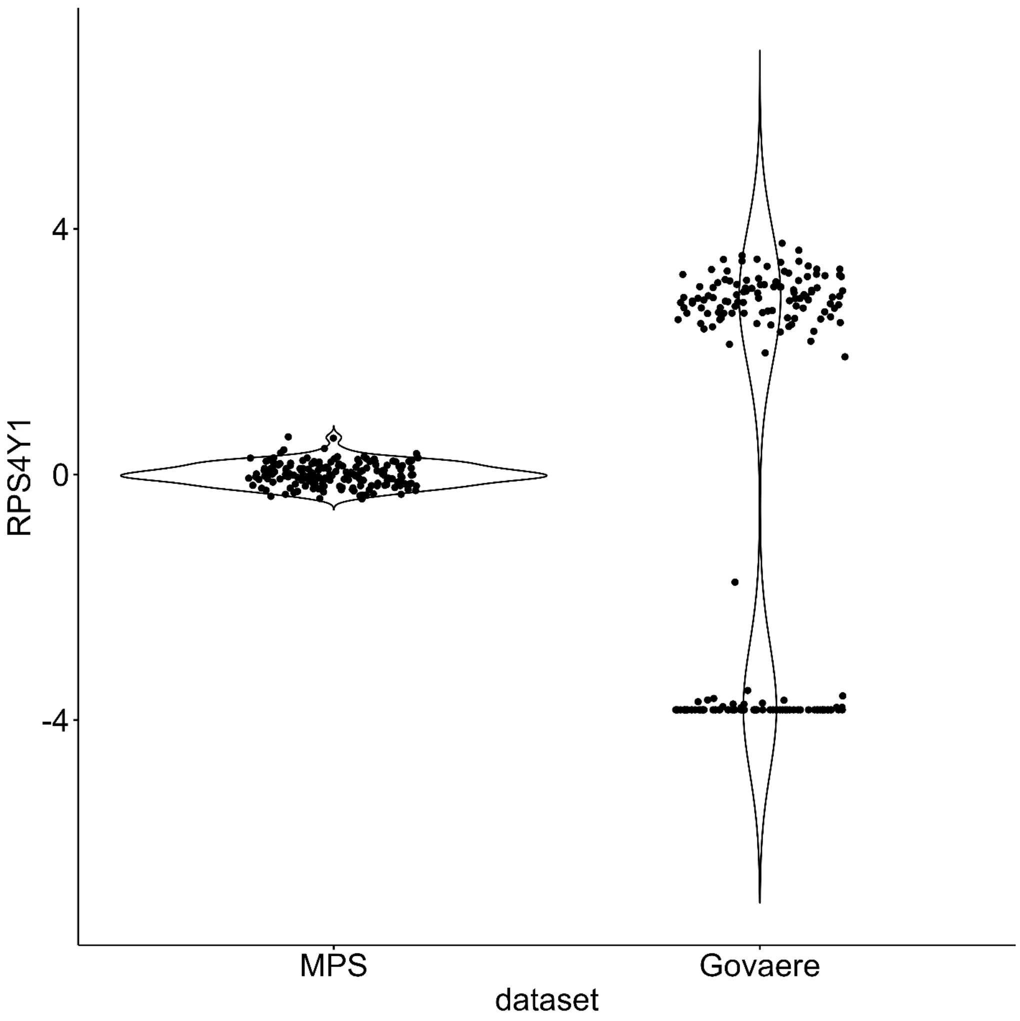


**Figure S13: Z-scored gene in the Y chromosome for the MPS and the clinical dataset.** The bimodal distribution in the clinical dataset is due to the inclusion of both male and female patients in the clinical study, while the lack of a bimodal distribution for the MPS indicates the use of only one sex.

### Supplementary Notes

We can show that the project-backproject step is equivalent to finding the best estimate of each human sample as a linear combination of all MPS samples. With the notation introduced in the methods section, $X_h$ is a matrix of normalized and centered human gene expression data (samples by genes), and $X_m$ is the normalized and centered MPS gene expression data. Consider the problem of estimating the gene expression profile of the $i$th human sample $X_{h,i}^T$ as a linear combination of the MPS gene expression data, where the observations are now the genes, and the features are the samples:

$$X_{h,i}^T \approx X_m^T \beta_i$$

With $\beta_i$ representing the weights of the linear combination. We can solve this problem through ordinary least-squares regression to find the weights. Since we can represent the MPS gene expression in terms of its principal component loadings $W_m$ and scores $S_m$ since by definition $X_m W_m = S_m \rightarrow X_m^T = W_m S_m^T$, we need to solve for:

$$X_{h,i}^T \approx W_m S_m^T \beta_i$$

Since the PCA loadings and scores are constant and depend only on the MPS dataset, we can redefine $\widetilde{\beta_i} = S_m^T \beta_i$ and find $\widetilde{\beta}_i$ directly, for which the least-squares solution is:

$$\widetilde{\beta_i} = \left(W_m^T W_m\right)^{-1} W_m^T X_{h,i}^T = W_m^T X_{h,i}^T$$

Where the product $W_m^T W_m = I$ because PCA loadings form an orthonormal basis by definition. The best estimate for a human sample $i$ is then:

$$\hat{X}_{h,i} = \left(W_m \widetilde{\beta_i}\right)^T = \left(W_m W_m^T X_{h,i}^T\right)^T = X_{h,i} W_m W_m^T$$

Stacking all the human samples we get:

$$\hat{X}_h = X_h W_m W_m^T$$

Which is equivalent to projecting human samples onto the MPS PCA space, and backprojecting to the original human gene expression space.
